# Replication-dead gammaherpesvirus vaccine protects against acute replication, reactivation from latency, and lethal challenge in mice

**DOI:** 10.1101/2023.09.26.559621

**Authors:** Wesley A. Bland, Shana Owens, Kyle McEvoy, Chad H. Hogan, Luciarita Boccuzzi, Varvara Kirillov, Camille Khairallah, Brian S. Sheridan, J. Craig Forrest, Laurie T. Krug

## Abstract

Gammaherpesviruses (GHVs) are oncogenic viruses that establish lifelong infections and are significant causes of human morbidity and mortality. While several vaccine strategies to limit GHV infection and disease are in development, there are no FDA-approved vaccines for human GHVs. As a new approach to gammaherpesvirus vaccination, we developed and tested a replication-dead virus (RDV) platform, using murine gammaherpesvirus 68 (MHV68), a well-established mouse model for gammaherpesvirus pathogenesis studies and preclinical therapeutic evaluations. We employed codon-shuffling-based complementation to generate revertant-free RDV lacking expression of the essential replication and transactivator protein (RTA) encoded by *ORF50* to arrest viral gene expression early after *de novo* infection. Inoculation with RDV-50.stop exposes the host to intact virion particles and leads to limited lytic gene expression in infected cells. Prime-boost vaccination of mice with RDV-50.stop elicited virus-specific neutralizing antibody and effector T cell responses in the lung and spleen. Vaccination with RDV-50.stop resulted in a near complete abolishment of virus replication in the lung 7 days post-challenge and virus reactivation from spleen 16 days post-challenge with WT MHV68. *Ifnar1^-/-^* mice, which lack the type I interferon receptor, exhibit severe disease upon infection with WT MHV68. RDV-50.stop vaccination of *Ifnar1^-/-^* mice prevented wasting and mortality upon challenge with WT MHV68. These results demonstrate that prime-boost vaccination with a GHV that is unable to undergo lytic replication offers protection against acute replication, reactivation, and severe disease upon WT virus challenge.

**Importance:** Gammaherpesviruses establish chronic infections that place a host at life-long risk for the development of lymphoproliferative disorders. While these viruses are endemic within the adult human population, there are currently no FDA-approved vaccines available to prevent acute infection or the later onset of lymphoproliferative disorders. We report here the use of a prime-boost vaccination strategy utilizing replication-dead virus strains to protect mice from acute disease following challenge with WT virus. We demonstrate that vaccination with replication-dead viruses is both safe and effective within immunocompetent and immunodeficient mice, and protects from viral replication, reactivation, and onset of severe disease.

## Introduction

Preventive vaccination against hepatitis B virus^1^ and high-risk human papillomaviruses^2^ has reduced the virus-associated cancer burden; however, an effective vaccine against the oncogenic gammaherpesviruses (GHVs) has not yet been deployed. GHVs include the human pathogens Epstein-Barr virus (EBV) and Kaposi sarcoma herpesvirus (KSHV). EBV causes infectious mononucleosis (IM) in adolescents and young adults during primary infection and is associated with the development of multiple sclerosis (MS)^3,4^. In addition, EBV is a cause of numerous lymphomas and epithelial-derived carcinomas of the oropharynx and gastrointestinal (GI) system^5,6^. KSHV is etiologically associated with lymphoproliferative diseases including primary effusion lymphoma (PEL) and a B cell variant of multicentric Castleman disease ^7^. KSHV is also the cause of Kaposi sarcoma, a tumor of presumed endothelial cell-origin that manifests in the skin, lungs, or GI tract^7,8^. Vaccines that block infection or lower viral burden are critical to develop as a means to reduce the incidence of GHV-associated diseases and cancers.

While knowledge of immune-dominant epitopes and correlates of immune protection for KSHV lag behind what is known for EBV, the need for a KSHV vaccine is paramount for at-risk populations^9^. People living with HIV (PWH) and transplant patients exhibit higher EBV and/or KSHV viral loads in circulating peripheral blood mononuclear cells (PBMCs) and are at increased risk for GHV-associated diseases and cancer, even when HIV is well controlled by anti-retroviral therapy^10,11,12^. KSHV is typically horizontally acquired during childhood in sub-Saharan Africa, and is a leading cause of morbidity and mortality, especially in geographical areas where malaria and HIV co-infections are endemic^13,14,15^.

GHVs, like all members of the herpesvirus family, display a biphasic infection cycle after entering a target cell. During productive lytic infection, a temporally regulated cascade of viral non-coding RNAs and gene products lead to the production of infectious particles capable of spreading to diverse cell types throughout the host. In contrast, the more immunologically silent latency phase is characterized by limited gene expression that promotes viability of the infected host cell and maintenance of the episomal viral genome^16^. Reactivation from latency to reinitiate the lytic cycle occurs intermittently in response to unknown stimuli and facilitates persistence in cellular reservoirs within the host and transmission to new hosts^17^. A vaccine that stimulates a broad humoral and cellular immune response against dozens of viral proteins in addition to the virion surface glycoproteins is hypothesized to have preventive and therapeutic potential.

Effective vaccine strategies have been developed against the human alphaherpesvirus varicella-zoster virus (VZV)^18,19^. Varivax, a live-attenuated VZV strain, protects against the onset of the childhood disease chickenpox following wild-type virus exposure. Notably, both wild-type and vaccine strains establish latent infection in dorsal root ganglia^20,21^. Reactivation and progression to shingles in elderly individuals infected with wild-type VZV is a painful illness with potential long-term complications including post-herpetic neuralgia^18^. The glycoprotein subunit-based vaccine Shingrix is a therapeutic vaccine widely administered to individuals over 50 years of age. Despite being administered after a person is colonized by VZV, Shingrix is remarkably effective at preventing shingles^22–27^. Thus, while it is typically considered ideal for potential vaccine candidates to prevent infection by generating sterilizing immunity, therapeutic vaccine strategies may also protect against the incidence of disease in individuals that harbor oncogenic GHVs.

To date, there are no FDA-approved vaccines that protect against human GHVs. Efforts focused on prophylactic vaccines to prevent virus infection have been challenged by the use of multiple glycoprotein complexes by GHVs to target different cell type-specific receptors. EBV glycoprotein gp350 binds to complement receptor CD21 on B cells^28^. In a phase 2 trial the administration of a gp350 subunit vaccine reduced infectious mononucleosis by 78% in participants, but had no impact on the rate at which the participants became infected with EBV^29,30^. Vaccines against EBV currently in trials present the gp350 subunit alone or in combination with glycoprotein subunits gH/gL/gp42 as a multivalent approach to broaden immunogenicity and block virus entry into B cells and epithelial cells^31^.

Due to the species-restricted nature of human GHVs, murine gammaherpesvirus 68 (MHV68), a GHV that infects murine rodents, is commonly used to study GHV pathogenesis and disease processes^32^. The MHV68 genome is colinear with KSHV, and the majority of ORFs encode structural and functional homologs of EBV and KSHV genes^33^. Importantly, like human GHVs, MHV68 infection establishes life-long infection in B cells and promotes diseases such as lymphoproliferative disorders in immunocompromised animals^34,35^. Over a span of three decades, viral pathogenic determinants of latency and replication, cellular reservoirs of infection, and key immune responses that control MHV68 infection in mice have been carefully delineated^35^. Live-attenuated viruses that replicate in the absence of latency protect against WT challenge^36–38^; transfer of serum and T cells from vaccinated mice prior to challenge protects against reactivation in recipient mice^36^. Thus, MHV68 infection of mice is a tractable small-animal pathogenesis system to evaluate the effectiveness of rationally designed vaccines and to identify immune correlates of protection against the pathogenic GHVs^39^.

Lytic replication and reactivation are controlled by the replication and transactivator (RTA) protein, a viral transcription factor encoded by *ORF50*^40,41^. In KSHV and MHV68, RTA promotes transcription of nearly all viral lytic genes^42^. Thus, RTA is essential for productive infection and lytic reactivation from latency. We recently developed a platform to produce high-titer, revertant-free stocks of a replication-dead virus (RDV) with a premature translational stop codon in *ORF50*^41,43^. Here, we test the efficacy of this forward-engineered infectious, yet replication-dead RTA-50.stop virus as a potential vaccine to stimulate an immune response that protects against wild-type MHV68 infection and virus-driven disease in mice. Immune correlates defined with this vaccine platform will inform KSHV vaccine design.

## Results

### Virus-specific adaptive immunity is induced by vaccination with RDV-50.stop

Insertion of a frameshift and translational stop codon in *ORF50* (ORF50.stop) renders MHV68 incapable of producing RTA, thereby generating RDV-50.stop^43^. RDV-50.stop particles undergo an abortive infection if not grown on a complementing cell line^43^. We expected that virion-associated proteins and limited RTA-independent viral gene expression would undergo antigen-presentation and promote virus-specific immune responses in vaccinated mice. Consistent with expectations, RNA-seq analysis of infected fibroblasts demonstrated that RDV-50.stop exhibits a global reduction in viral gene expression when compared to WT MHV68 (Fig. 1a). Although median expression levels were reduced ∼300-fold compared to WT infection, transcripts for ORFs associated with latency (*M3*, *M4* and *ORF73*/LANA) and lytic replication that are also immunodominant epitopes (*ORF6* and *ORF61*) were detected upon infection with RDV-50.stop. Given that RDV-50.stop infection is self-limiting and viral gene expression is sharply curtailed, we hypothesized that vaccination of mice using RDV-50.stop would require multiple doses to generate a potent MHV68-specific immune response.

**Fig 1.**
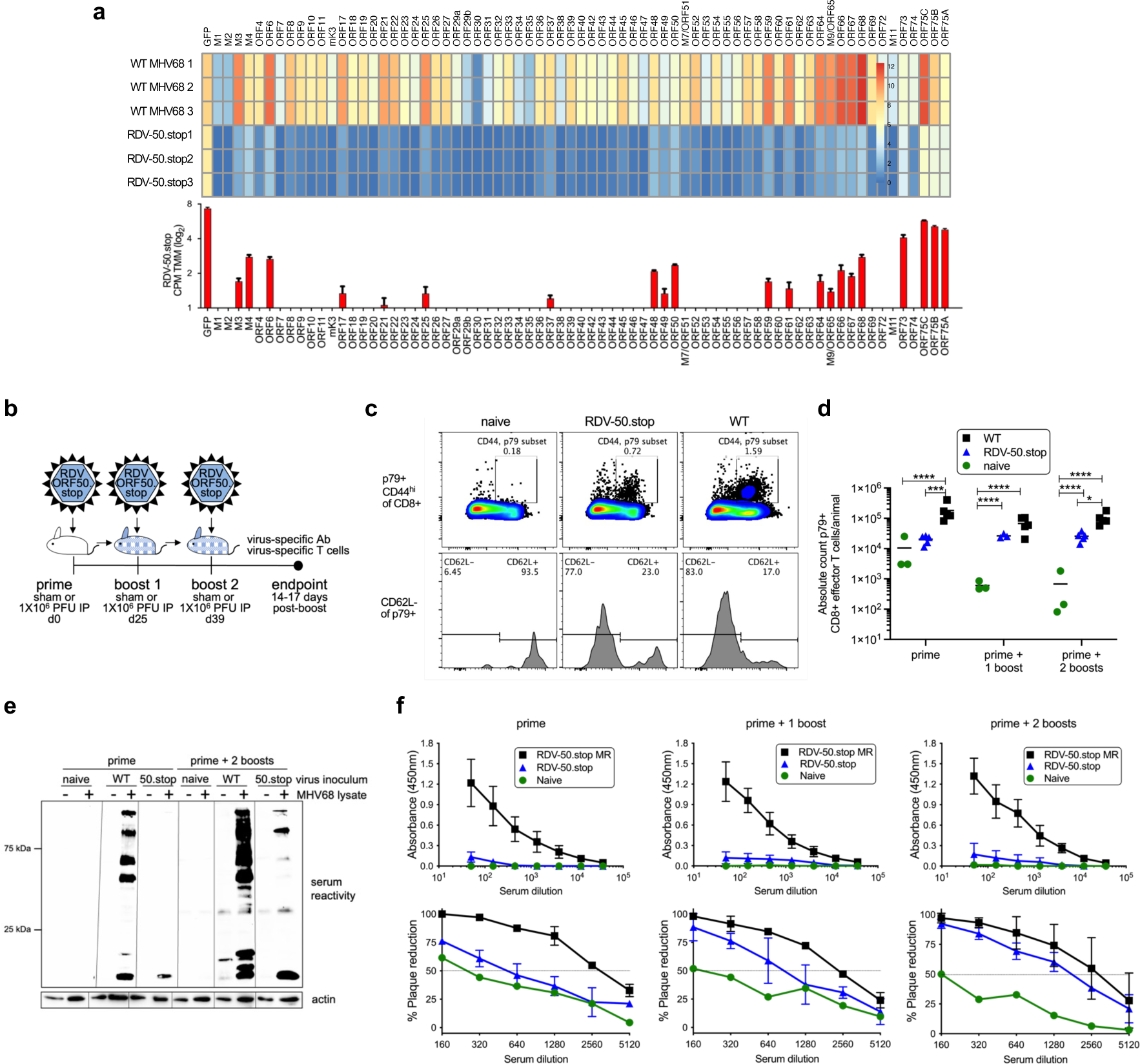
A replication-dead recombinant MHV68 generates virus-specific immune responses upon a prime-boost regimen in C57BL/6 mice. **a** Top panel, RNAseq profile of viral gene expression in murine fibroblasts 18 hpi with RDV-50.stop and WT MHV68 (MOI 3) in biological triplicate. Scale represents log_2_ CPM TMM value for non-overlapping ORF regions. Lower panel, bars (mean +/-SD) represent ORF gene expression upon RDV-50.stop infections. **b** Schematic of prime-boost strategy to examine the immune response of C57BL/6 mice to RDV-50.stop and WT control virus infections. Naïve mice were age-matched, non-vaccinated controls. **c** Virus-specific effector CD8-T cell response based on p79 tetramer+ of CD44^hi^CD62L^-^ CD8 T cells. Representative gating strategy from naïve mice or mice infected with RDV-50.stop of WT virus at d39. **d** Total p79-reactive CD8 T cells per spleen of individual mice after initial prime and sequential boosts. Symbols represent individual mice (N=3-5) and bars are mean values; *, p<0.05; **, p<0.002; ***, p<0.0002; ****, p<0.0001 in Tukey’s multiple comparisons post-test of two-way ANOVA. **e** Immunoblot analysis of infected cell lysates testing reactivity of sera from immunized mice after initial prime or prime followed by two boosts. Lines indicate blot strips incubated with sera from indicated animals and then reassembled for imaging. **f** Top, panel defining virus-specific IgG in the sera of mice after prime and post-boost. Symbols represent individual mice (N=5); bars are mean values +/-SD. Below, neutralizing antibodies were evaluated by determining the serum dilution that led to a 50% reduction in plaque formation. Symbols represent individual mice (N=3); bars are mean values +/-SD.

We tested this hypothesis using the common prime-boost strategy of vaccination. C57BL/6 mice were inoculated intraperitoneally (i.p.) with 10^6^ PFU of RDV-50.stop (day 0). One experimental group (the prime-boost group) received one boost at day 25. A separate set of animals received two boosts (prime-boost-boost), one at day 25 and another on day 39 after the initial inoculation (Fig. 1b). We evaluated virus-specific CD8 T cell and humoral immune responses 14 days after the final RDV-50.stop dose in comparison to mice infected with WT MHV68. CD8 T cells that are specific for lytic antigens restrict virus replication through cytolytic activity and the secretion of antiviral cytokines such as interferon-γ^44–46^. Virus-specific effector CD8 T cells (CD8 T^eff^) were identified by flow cytometry as CD19^-^, CD3^+^, CD8^+^, CD44^hi^, CD62L^+^ and tetramer-positive for the p79 immunodominant epitope derived from ORF61 (Fig. 1c). As expected, although generally lower than numbers induced by WT virus infection, virus-specific CD8 T^eff^ were significantly elevated after vaccination with RDV-50.stop, with comparable detection following either single or double boosting regimens (Fig. 1d & Supplementary Fig. S1). These findings demonstrate that sufficient viral antigen is available and presented in a manner following vaccination with a replication-dead GHV to elicit virus-specific CD8 T^eff^ responses in mice.

Infection with MHV68 stimulates a potent polyclonal antibody response that is maintained over-time^47–49^. Passive serum transfer suggests that the antibodies generated during MHV68 infection are neutralizing and contribute to control of WT virus infection^50,36^. To evaluate antibody responses to RDV-50.stop, we first compared lytic viral antigen reactivity in immunoblot analyses following priming alone, double boosting, and WT MHV68 infection. While a single dose of RDV-50.stop elicited minimal reactivity, the three-dose vaccination strategy stimulated a potent response, with antibodies recognizing many viral antigens that were also recognized by sera from animals infected with WT MHV68 (Fig. 1e). The total MHV68-specific IgG was only slightly elevated over sham-vaccination controls by ELISA, even following three doses of RDV-50.stop, and were substantially lower than IgG levels produced by WT MHV68 infection of mice (Fig. 1f, upper panel). However, 50% plaque-reduction neutralizing titer (PRNT50) assays revealed that the relatively low quantity of virus-specific IgG produced by vaccination with RDV-50.stop was capable of blocking MHV68 infection (Fig. 1f, lower panel). These findings indicate that vaccination with RDV-50.stop stimulates potent and effective neutralizing antibody production, despite eliciting MHV68-directed IgG to much lower levels than WT virus infection. Together, these data demonstrate that a prime-boost vaccination strategy using a replication-dead GHV elicits virus-specific CD8 T^eff^ and neutralizing antibody responses *in vivo*.

### Vaccination with RDV-50.stop protects mice from WT challenge

Since vaccination with RDV-50.stop promoted MHV68-directed cellular and humoral immune responses, we sought to determine if the vaccine response was protective *in vivo*. C57BL/6 mice were vaccinated with RDV-50.stop as described above or using PBS as a sham control. 14-17 days after the final dose, mice were challenged intranasally with 1000 PFU of WT MHV68 (Fig. 2a). Plaque assays performed on homogenized lung tissue 7 days post-infection revealed significantly lower viral titers in lungs of vaccinated animals, with mice that received either one or two boosts exhibiting a three-log reduction in viral titers compared to sham-vaccinated controls (Fig. 2b). These data demonstrate that vaccination with RDV-50.stop potently inhibits WT virus replication at a mucosal barrier.

**Fig 2.**
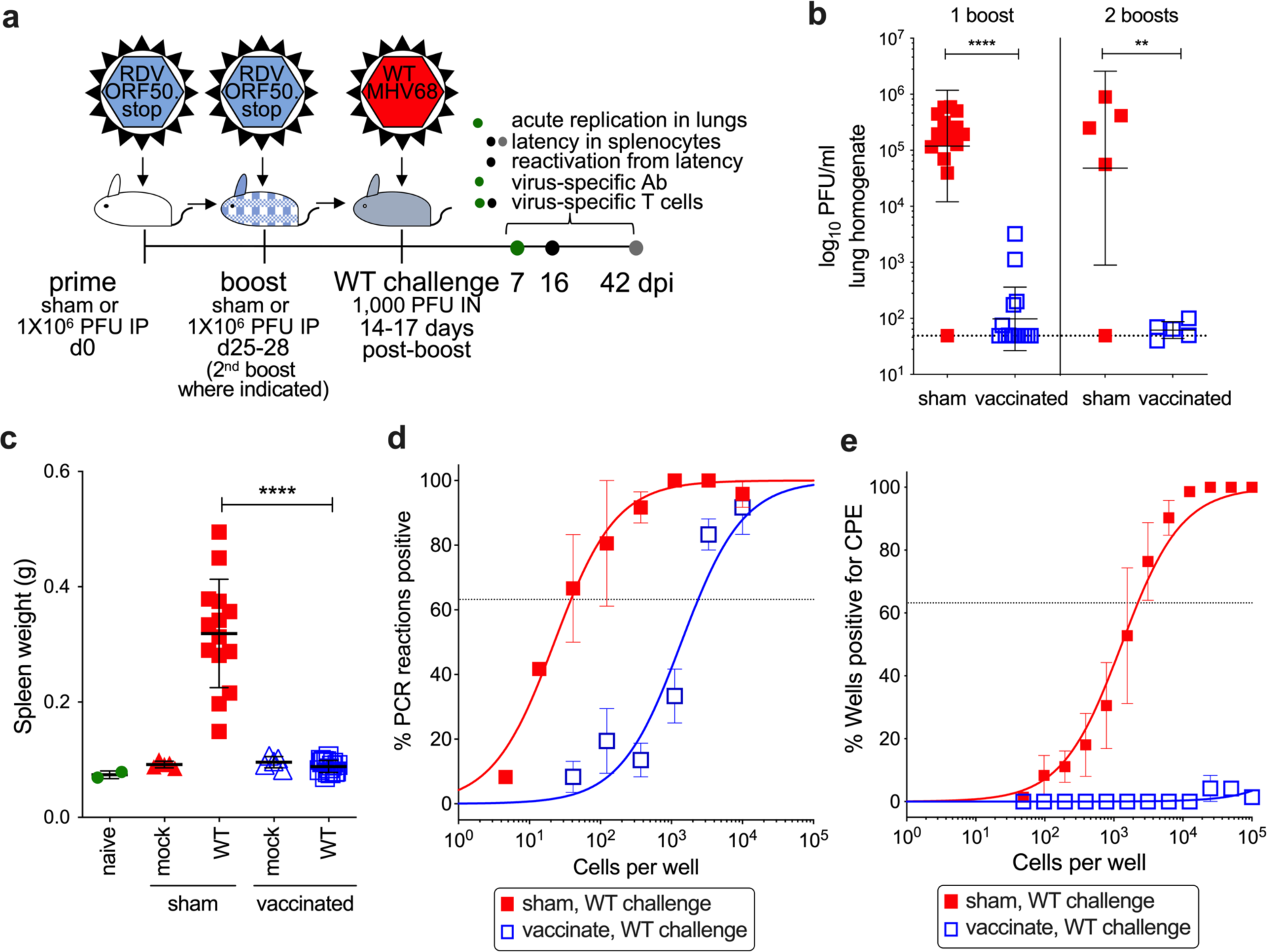
Vaccination with RDV-RDV-50.stop reduces acute replication, splenomegaly, latency and reactivation of wild-type MHV68 upon challenge in C57BL/6 mice. **a** C57BL/6 mice were either sham-vaccinated or vaccinated and then boosted once **b-e** or twice b with 1×10^6^ PFU RDV-50.stop followed by challenge with 1×10^3^ PFU WT MHV68 at 14-17 d post-boost. **b** Acute replication at 7 dpi determined by measuring infectious particles per ml lung homogenate. Symbols denote individual mice from one (2 boosts) or three (1 boost) independent experiments for infected animals (N=5 mice per experiment); bars and whiskers represent mean+/-SD. **, p<0.01; ****, p<0.0001 in one-tailed unpaired T test of log-transformed values. **c** Splenomegaly determined by spleen weights. Symbols denote individual mice (N=2-15); bars and whiskers represent mean +/-SD. ****, p<0.0001 in Tukey’s test of one-way ANOVA between the sham and vaccinated groups upon WT challenge at 16 dpi. **d** The frequency of latency determined by limiting dilution nested PCR of intact splenocytes for the viral genome at day 16 post WT challenge. **e** The frequency of explant reactivation determined by limiting dilution coculture of intact viable splenocytes on a monolater of primary MEFs 16 days post WT challenge. Disrupted splenocytes plated in parallel did not detect preformed infectious virus in the vaccinated animals. **d-e** Symbols denote the average of three experiments with five mice per experiment; error bars represent SEM.

Since latent GHV infection is associated with numerous malignancies, a major goal in the development of GHV vaccines is to reduce latent infection and reactivation. We therefore sought to determine the impact of RDV vaccination on MHV68 latency. MHV68 reaches the splenic lymphoid tissue after viral transit through the lymph node and hematogenous dissemination ^51^. An increase in spleen weight reflects MHV68 colonization of the spleen that occurs in an infectious mononucleosis-like syndrome during MHV68 latency establishment^52^. Infection of sham-vaccinated animals with WT MHV68 resulted in a three-fold increase in spleen weight relative to naive mice at 16 dpi (Fig. 2c). While vaccination alone did not cause an increase in spleen weight, vaccination with RDV-50.stop prior to WT virus challenge prevented infection-related splenomegaly. In limiting-dilution PCR analyses to measure the frequency of splenocytes that harbor MHV68 genomes, we found that vaccination prior to WT virus challenge infection resulted in a profound reduction in latently infected cells from approximately 1 in 38 splenocytes containing viral genomes in sham-vaccinated animals to 1 in 2380 after vaccination (Fig. 2d). When we extended this analysis to 42 dpi, we observed overlapping frequencies of MHV68 genome-positive splenocytes in sham-vaccinated and vaccinated animals (Supplementary Fig. S2a).

Latent GHVs undergo periodic reactivation which facilitates virus persistence and transmission and is also thought to contribute to virus-driven oncogenesis and inflammatory diseases^6,53,54^. To determine the effect of vaccination with RDV-50.stop on MHV68 reactivation from latency, we performed serial dilutions of explanted splenocytes from vaccinated and sham-vaccinated animals on an indicator monolayer and evaluated cytopathic effects. While splenocytes from sham-vaccinated mice exhibited a reactivation frequency of approximately 1 in 2000 splenocytes, reactivation frequencies were below the limit of detection after challenging vaccinated animals with WT MHV68 (Fig. 2e). These data suggest that immunity induced by vaccination with RDV-50.stop inhibits MHV68 reactivation.

RTA expression is not essential to establish latent infection of B cells in vivo^55^. LANA, the viral episomal maintenance protein encoded by *ORF73*, and other viral latency-associated genes were expressed in the absence of RTA (Fig. 1b), which led us to hypothesize that RDV-50.stop was competent for latency establishment in the spleen. To differentiate the RDV-50.stop vaccine virus from WT challenge virus, we performed semi-quantitative PCR analyses using RDV-50.stop specific primer pairs to determine the relative percentage of vaccine-derived virus present in spleens after WT challenge. As expected, each group of animals infected with a single strain of virus (either RDV-50.stop or WT MHV68) was positive only for the respective virus they received (Supplementary Fig. S2b). In contrast, DNA from spleens of vaccinated mice that were challenged with WT virus contained a mixture of viral genomes. In a standardized quantity of spleen DNA, only 13.9% of viral DNA amplified was from the vaccine strain. This indicates that, while lytic replication and latency establishment by WT MHV68 is potently restricted by prime-boost vaccination with RDV-50.stop, it does not elicit sterilizing immunity against WT virus infection.

### Virus-specific immune response correlates with protection from WT challenge

Identification of immune correlates of protection is an important aspect of developing an effective vaccine for GHVs and other viruses. We therefore determined the effect RDV-50.stop vaccination had on immune activation following WT MHV68 infection. In analyses of T cell responses during the acute phase of WT virus challenge, CD8 T cells specific for the MHV68 p79 and p56 immunodominant epitopes were detected in the lungs, the primary site of infection after IN inoculation, of vaccinated mice 7 days after WT challenge (Fig 3a-b). In spleens, CD8 T cells specific for p79 (Fig. 3c-d) and p56 (Supplementary Fig. S3a) were abundant in vaccinated mice, with or without infection. Further immunophenotyping revealed that over 50% of virus-specific CD44^+^ CD8 T cells in vaccinated mice were KLRG1^+^ CD127^-^, consistent with a short-lived effector cell (SLEC) phenotype (Fig. 3c,e and Supplementary Fig. S3b). Analysis of KLRG1^-^ CD127^+^ memory precursor effector CD8 T cells (MPEC) by CD62L identified a higher proportion of CD62L^-^ MPEC compared to CD62L^+^ MPEC in vaccinated mice compared to their sham-vaccinated counterparts (Fig 3c,f-g and Supplementary Fig. S3c-d). Stimulation of splenic CD8 T cells with p79 (Fig. 3h) and p56 (Fig. 3i) peptides led to increased production of antiviral effector cytokines TNFα and IFNγ in vaccinated mice (Fig. 3j), demonstrating that antigen-specific cells were present. Virus-specific neutralizing antibodies were also detected in RDV-50.stop vaccinated mice 7 days post-challenge (Supplementary Fig. S3e-f). Taken together, prime-boost vaccination with RDV-50.stop elicited virus-specific CD8 T cells that are poised to respond to WT challenge during the acute phase of infection. Thus, neutralizing antibody and virus-specific CD8 T_eff_ responses correlate with the strong protection against WT replication and reactivation from latency that is afforded to vaccinated mice (Fig. 2).

**Fig 3.**
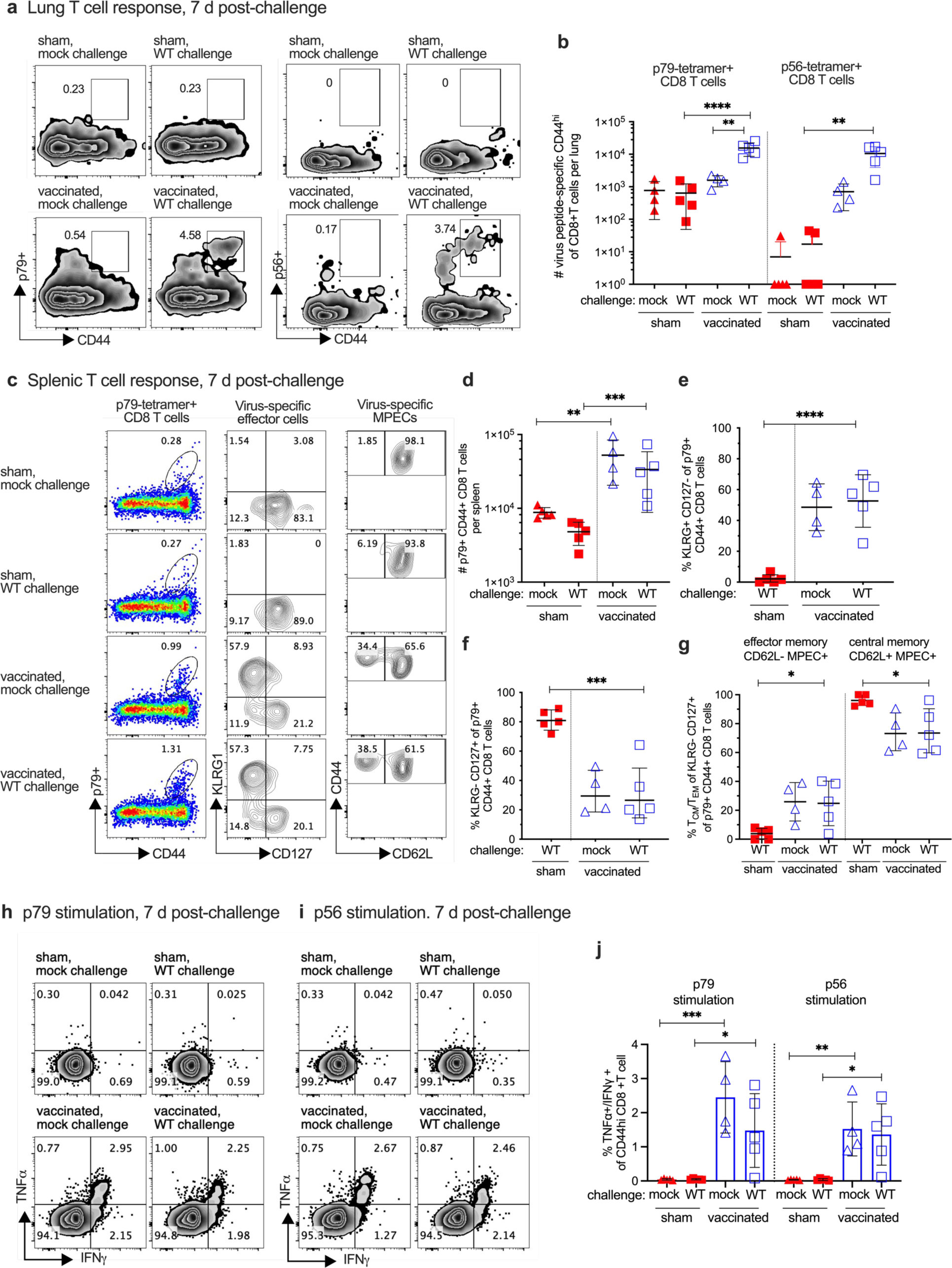
Evaluation of T cell response to MHV68 in the lungs and spleens of vaccinated mice at seven days post-challenge with WT virus. C57BL/6 mice were either sham-vaccinated or vaccinated twice (prime+boost) with 1×10^6^ PFU RDV-50.stop followed by mock challenge or challenge with 1×10^3^ PFU WT MHV68 at 15 d post-boost and analyzed 7 d post-challenge. **a** Flow cytometric gating strategy to determine the frequency of CD44^hi^ CD8 T cells in the lungs that were reactive with viral p79 or p56 epitopes. **b** Total p79-or p56-tetramer CD8 T cells per spleen of individual mice after initial prime and sequential boost. **c** Left column, flow cytometric gating strategy to determine the frequency of CD44^hi^ CD8 T cells in the spleens that were reactive with viral p79. Middle column, p79-tetramer+ CD8 T cells were further analyzed for markers of short-lived effector cell (SLEC, KLRG^+^CD127^-^) and memory precursor effector cell subsets (MPEC, KLRG^-^CD127^+^). Right column, MPECs were further delineated into CD62L^-^ effector and CD62L^+^ central MPECs. **d** Total p79-tetramer+ CD8 T cells per spleen of individual mice with the indicated vaccination and challenge regimen. The percentage of p79-reactive CD8 T cells that were SLECs **e**, MPECs **f,** and effector vs memory MPECs **g** were enumerated based on the gating strategy for surface markers in **c**. Intracellular cytokine levels of effector cytokines TNFα and IFNγ were examined 6 hrs after stimulation with **h** p79 and **i** p56 peptides. **j** Percentage of CD44^hi^ CD8 T cells producing both TNFα and IFNγ. For each graph, symbols represent individual mice, (N=4-5); bars and whiskers are mean +/-SD values.*, p<0.05; **, p<0.05; ***, p<0.001; ****, p<0.0001 in Sidak’s multiple comparisons test of one-way ANOVA between the indicated groups.

The CD8 T cell response was next compared between the sham-vaccinated and RDV-50.stop vaccinated mice at 16 day post-WT challenge, a timepoint of peak virus latency in the early chronic phase of WT infection. A potent virus-specific CD8 T cell response was observed at 16 dpi in the lungs of sham-vaccinated mice that surpassed the levels detected in the vaccinated mice upon challenge (Fig. 4a-c), consistent with the typical kinetics of the CD8 T cell response that arrests the WT virus replication observed at 7 dpi (Fig. 2b). A similar observation was made for the spleen at 16 dpi. Virus-specific CD8 T cells in sham-vaccinated animals exceeded those in the vaccinated mice, and these T cells had an increased SLEC and CD62L^-^ MPEC phenotype at 16 dpi, for both p79 (Fig. 4d-g) and p56 (Supplementary Fig. S4a-d) epitopes. Antiviral TNFα and IFNγ effector cytokines correlated with this increased effector phenotype in spleens of WT infected mice that were not vaccinated compared to their RDV-50.stop vaccinated counterparts, with or without challenge (Fig 4h-j). In summary, vaccinated mice challenged with WT virus exhibited a reduced CD8 T cell response compared to their non-vaccinated counterparts at 16 dpi. This observation is consistent with a strong pre-existing immune response in the lungs and spleen afforded by RDV-50.stop vaccination (Fig. 3) that limits viral replication (Fig.2). In contrast, sham-vaccinated mice are unable to control viral replication and elicit a larger CD8 T cell response with delayed kinetics (Fig. 1).

**Fig 4.**
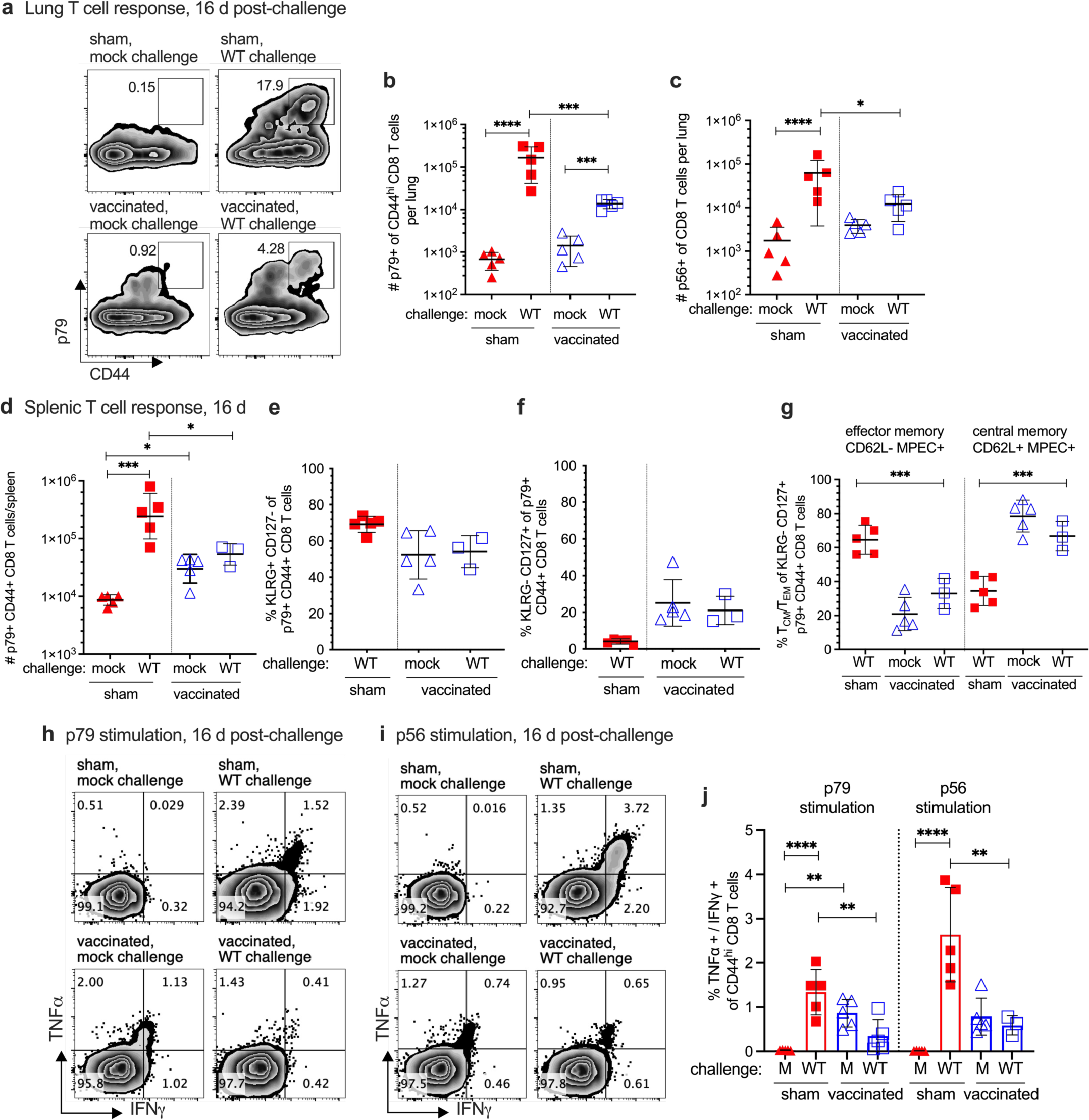
Evaluation of T cell response to MHV68 in the lungs and spleens of vaccinated mice at sixteen days post-challenge with WT virus. C57BL/6 mice were either sham-vaccinated or vaccinated twice (prime+boost) with 1×10^6^ PFU RDV-50.stop followed by mock challenge or challenge with 1×10^3^ PFU WT MHV68 at 15 days post-boost. **a** Flow cytometric gating strategy to determine the frequency of CD44^+^ CD8 T cells in the lungs of individual mice with the indicated vaccination and challenge regimen that were reactive with viral p79 at 16 days post challenge. **b-c** Total p79-or p56-tetramer+ CD8 T cells per lung of individual mice after initial prime and sequential boost. **d** Total p79-tetramer+ CD8 T cells per spleen of individual mice after initial prime and sequential boosts. Percentage of p79-tetramer+ CD8 T cells with markers of **e** short-lived effector cell (SLEC, KLRG^+^CD127^-^) and **f** memory precursor effector cell subsets (MPEC, KLRG^-^ CD127^+^). **g** MPECs were further delineated into CD62L^-^ effector and CD62L^+^ central MPECs for p79-tetramer+ CD8 T cells. Intracellular cytokine levels of effector cytokines TNFα and IFNγ were examined 6 hrs after stimulation with **h** p79 and **i** p56 peptides. **j** Percentage of CD44^hi^ CD8 T cells producing both TNFα and IFNγ in response to viral peptide stimulation. For each graph, symbols represent individual mice, (N=3-5); bars and whiskers are mean +/-SD values. *, p<0.05; **, p<0.05; ***, p<0.001; ****, p<0.0001 in Sidak’s multiple comparisons test of one-way ANOVA between the indicated groups.

### Durable protection with RDV-50.stop infection

To test the durability of RDV-50.stop vaccine-induced immunity, we evaluated the impact of vaccination on WT virus challenge 90 days after completion of the prime-boost regimen (Fig. 5a). In vaccinated mice, viral titers in lungs were undetectable by plaque assay in four of five animals at 7 days post-challenge (Fig. 5b). This correlated with detection of virus-specific, neutralizing antibodies (Fig. 5c-d) and the presence of virus-specific CD8 T cells in the lungs of vaccinated mice both before and after challenge (Fig. 5e). Moreover, CD8 T cells recognized by the p56 MHCI tetramer in spleens of vaccinated mice bore markers of effector cells (Fig. 5f-g) and produced antiviral cytokines TNFα and IFNγ upon peptide stimulation (Fig. 5h).

**Fig 5.**
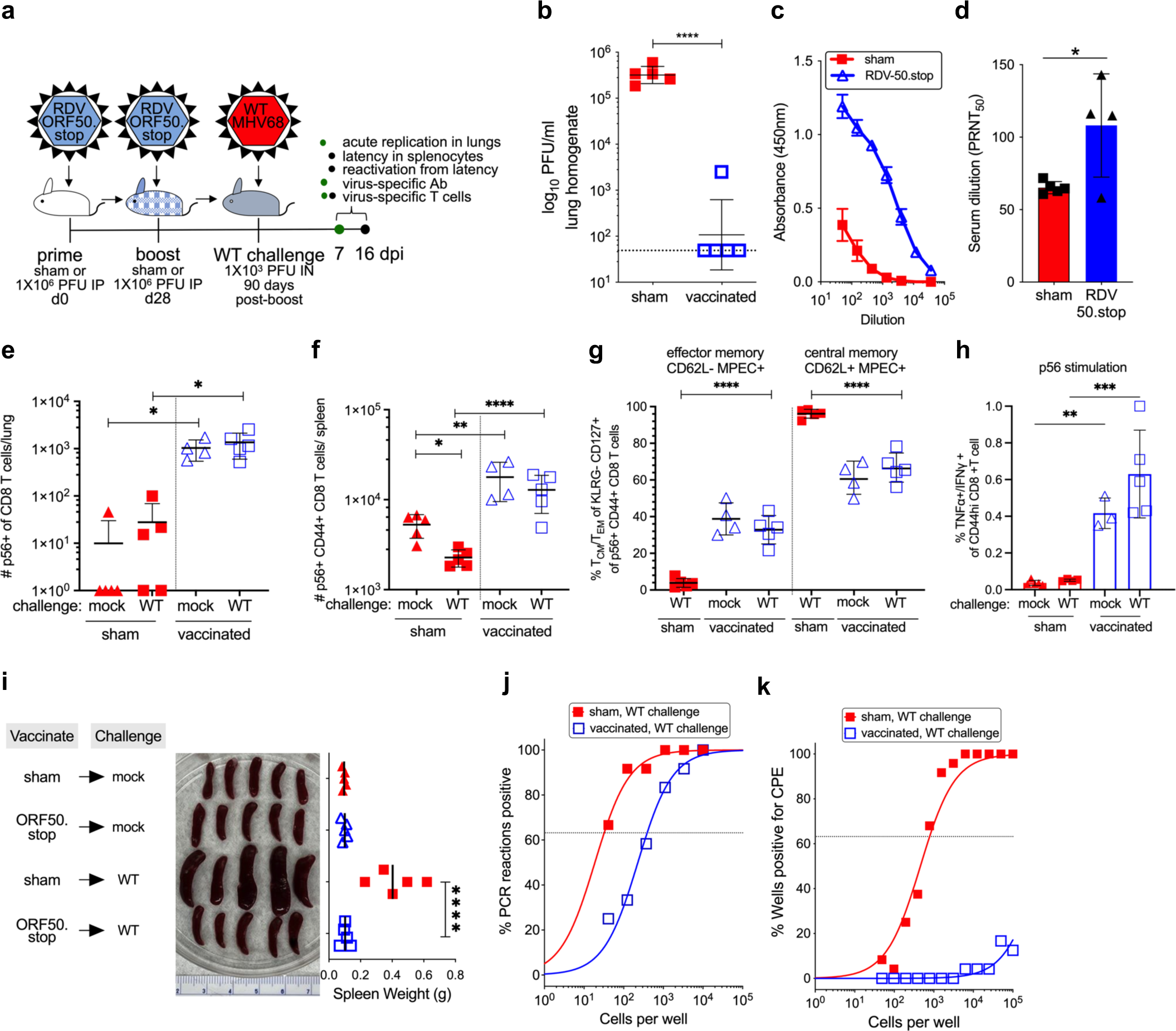
Vaccination with RDV-50.stop leads to durable protection against wild-type MHV68 challenge. **a** C57BL/6 mice were either sham-vaccinated or vaccinated twice (prime+boost) with 1×10^6^ PFU RDV-50.stop followed by challenge with 1×10^3^ PFU WT MHV68 at 90 days post-boost. **b** Acute replication at 7 dpi determined by measuring infectious particles per ml lung homogenate. Symbols denote individual mice (N=5); bars and whiskers represent mean+/-SD. ****, p<0.0001 in one-tailed unpaired T test of log-transformed values. **c** Virus-specific IgG from sham or RDV-50.stop vaccinated mice at 7 d post challenge measured by ELISA. Symbols denote individual mice; bars and whiskers represent mean+/-SD. *, p<0.05 in two-tailed unpaired T test. **d** Virus neutralization in serum as determined by a plaque reduction assay. The plaque reduction neutralization titer 50 (PRNT_50_) value is the dilution of serum to reach 50% neutralization of plaques. **e-f** Total p56-tetramer+ CD8 T cells per lung e or spleen f of individual mice. **g** MPECs were delineated into CD62L^-^ effector and CD62L^+^ central MPECs for p56-tetramer+ CD44^+^ CD8 T cells. **h** Percentage of CD44^hi^ CD8 T cells producing both TNFα and IFNγ in response to viral p56 peptide stimulation. For **d-h**, symbols represent individual mice, (N=3-5); bars and whiskers are mean +/-SD values. *, p<0.05; **, p<0.05; ***, p<0.001; ****, p<0.0001 in Dunn’s **e** or Sidak’s **f-h** multiple comparisons test of one-way ANOVA between the indicated groups. **i** Splenomegaly visualized for whole spleens and quantitated by spleen weights. For, symbols represent individual mice, (N=5); bars and whiskers represent mean +/-SD. ****, p<0.0001 in Tukey’s test of one-way ANOVA. **j** The frequency of latency determined by limiting dilution nested PCR of intact splenocytes for the viral genome. **k** The frequency of explant reactivation determined by limiting dilution coculture of intact viable splenocytes on a monolater of primary MEFs. Disrupted splenocytes plated in parallel did not detect preformed infectious virus in the vaccinated animals. Error bars represent standard error of the means. Symbols denote the average of one experiment with five mice per experiment.

With regard to durability of protection against WT MHV68 latency establishment, splenomegaly was reduced by four-fold in RDV-50.stop vaccinated mice compared to sham-vaccinated mice on day 16 after intranasal challenge with WT virus (Fig. 5i). Congruent with decreased splenomegaly, vaccination also resulted in a 10-fold reduction in the frequency of viral genome-positive cells in comparison to non-vaccinated controls (Fig. 5j), and virus reactivation was well below the threshold of accurate quantitation, reduced by at least three orders of magnitude (Fig. 5k). By 16 days post WT challenge, the number of virus-reactive CD8 T cells in lungs (Supplementary Fig. S5a-b) and spleens (Supplementary Fig. S5c,g) were lower in vaccinated compared to sham-vaccinated animals, but they had higher frequencies of MPEC (Supplementary Fig. S5d-e,i) and CD62L^-^ MPEC CD8 T cells (Supplementary Fig. S5f,j) in sham-vaccinated mice compared to RDV-50.stop vaccinated mice, with or without challenge. This suggests that either the anamnestic response to re-infection in vaccinated animals requires less activation and expansion to be effective or that local immune control in the lung reduces the general need for immune expansion to control MHV68 infection. Overall, these data indicate that the protective immune response generated by vaccination with RDV-50.stop is durable and protective up to 3 months post-boost.

### Vaccination with RDV-50.stop protects mice lacking type I interferon responses from severe disease

GHV infection can lead to a diverse array of lymphoproliferative disorders, inflammatory diseases, and even death in hosts with weakened adaptive and innate immune systems^56^. Type I interferon signaling is required to control lytic replication and reactivation of MHV68 from latency^57,58^. Mice lacking *Ifnar11* are incapable of responding to interferon-ɑ/β, leading to an increase in susceptibility to severe disease, acute viral replication, and mortality when infected with MHV68^59^. To determine if RDV-50.stop vaccination was protective in a host that is highly susceptible to severe disease, we vaccinated mice lacking the type I interferon receptor common alpha subunit (*Ifnar1^-/-^)*.

*Ifnar1^-/-^* mice were sham-vaccinated or vaccinated according to the previously established timeline (Fig. 6a). Similar to WT C57BL/6 mice, vaccination of *Ifnar1^-/-^* mice with RDV-50.stop resulted in an increase in virus-specific, neutralizing antibody production (Supplementary Fig. S6). Two weeks after completing the vaccination regimen, *Ifnar11^-/-^* mice were challenged intranasally with 2×10^6^ PFU of WT MHV68, a dose previously determined to be lethal in *Ifnar1^-/-^* mice on a C57BL/6 background^60^, and monitored for signs of disease for 6 weeks after challenge. Mice were weighed daily for 42 days and assessed for signs of distress. Sham-vaccinated animals began losing weight on day 5 post-challenge, with 90% of animals losing greater than 10% body weight by day 10 post-challenge (Fig. 6b and Supplementary Fig. S7). By 11 days post-challenge, 6 out of 10 sham-vaccinated *Ifnar1^-/^*^-^ reached humane endpoint criteria (Fig. 6c). In contrast, none of the vaccinated *Ifnar1^-/-^*mice showed signs of disease; weights remained stable, and animals remained mobile and active throughout the course of the study (Fig. 6b,c and Supplementary Fig. S7).

**Fig 6.**
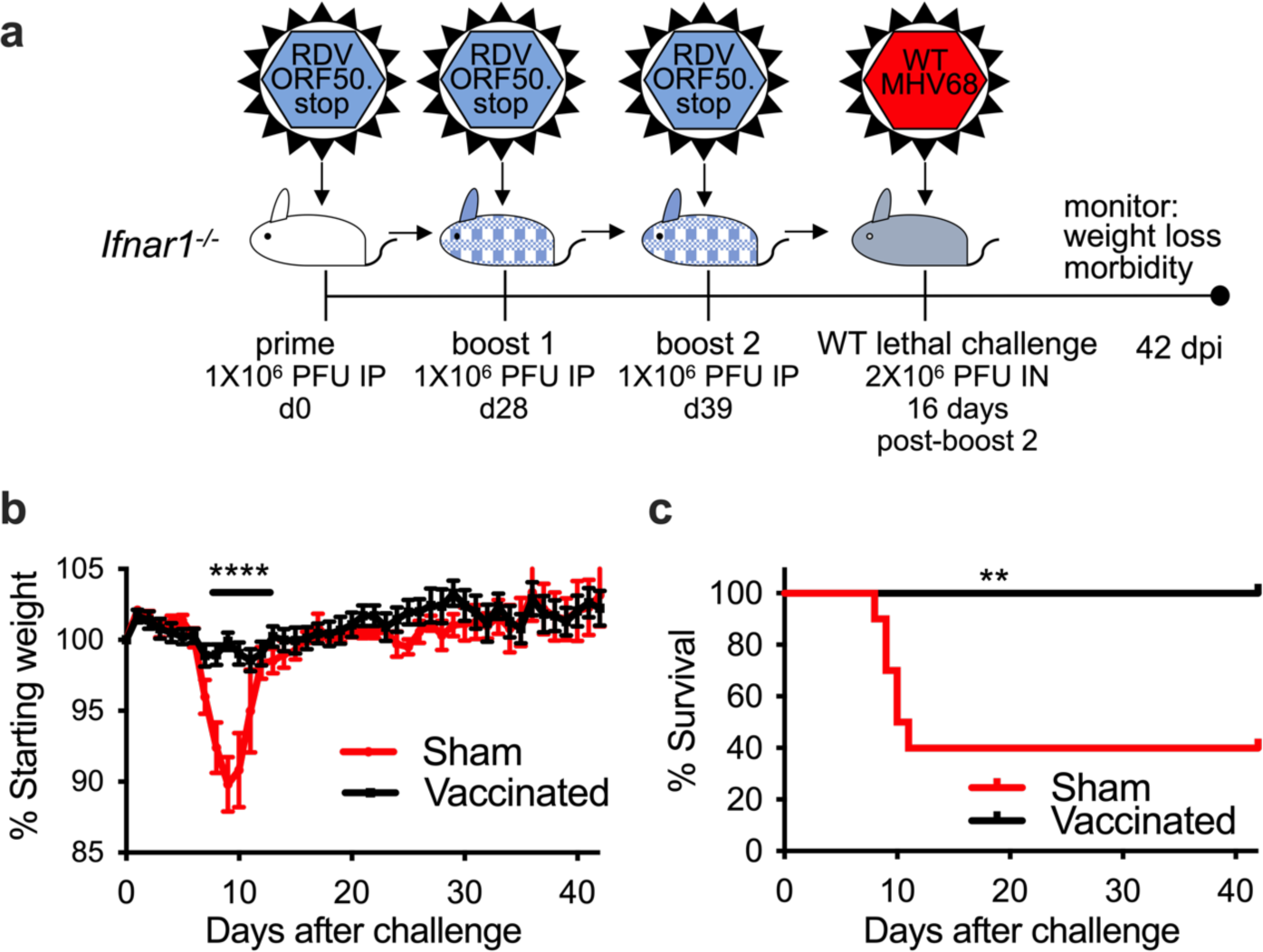
Vaccination with RDV-50.stop protects mice susceptible to severe disease from weight-loss and disease. **a** Groups of 10 C57BL/6 *Ifnar1^-/-^* mice were either sham-vaccinated or vaccinated with 3 doses of 1X 10^6^ PFU RDV-50.stop MHV68 and challenged with a lethal dose of 2×10^6^ PFU WT MHV68. Sham-vaccinated and vaccinated mice were weighed daily for 42 days to measure disease progression. **b** Daily weights were averaged to determine protective effect of the vaccinated group. **c** Survival curve for sham-vaccinated and vaccinated *Ifnar1^-/-^*mice. Symbols represent individual mice; error bars represent SEM. ****, p<0.0001 in a two-tailed student’s t-test; **, p<0.01 in a logrank test.

Consistent with effects of vaccination observed for WT mice, vaccination with RDV-50.stop MHV68 resulted in a significant decrease in the frequency of splenocytes harboring viral genomes. For surviving sham-vaccinated mice, approximately 1 in 100 splenocytes were viral-genome positive, compared to 1 in 3500 splenocytes in vaccinated animals (Supplementary Fig. S8a). When the source of viral genomes was evaluated by genotyping PCR, 14% of the virus present was vaccine-derived (Supplementary Fig. S8b), which further suggests that vaccination with RDV-50.stop, while clearly protective against severe disease, does not offer sterilizing immunity against latent colonization by a WT virus.

## Discussion

In this study, we demonstrate the potential for a replication-dead gammaherpesvirus to elicit a protective immune response and protect from WT replication, latency establishment, and disease *in vivo*. RDV-50.stop functions as a hybrid vaccine platform, capitalizing on the complex antigenicity of the virus particle itself combined with its ability to infect antigen-presenting cells to permit antigen presentation of virion-associated proteins and the limited genes expressed from the delivered viral genome.

Cellular immune responses are understudied in the context of human GHV vaccinology. MuMT mice, which lack the capacity to form B cells, are capable of controlling acute MHV68 infection due to the cellular response to infection^61^. Mice lacking either CD4 or CD8 T cells are less capable of controlling acute viral replication and reactivation from latency, resulting in increased susceptibility to severe disease and lymphoproliferation^44,62–66^. Here, humoral and cellular mediated immune responses of vaccinated mice were monitored prior to and upon WT challenge. The neutralizing titer of serum and effector T cells specific to known immunodominant epitopes from lytic proteins increased following prime-boost dosing of WT C57BL/6 mice with RDV-50.stop. Antibody functions likely extend beyond neutralization and FcR-mediated enchacement of entry should be evaluated. Clearance of acute replication of MHV68 in the lungs requires CD8, but not CD4 T cells^63,67^. In the lungs and spleens of vaccinated mice, there is an effector CD8 T cell population specific for the major lytic viral antigens p79, an epitope present within the viral large ribonucleotide reductase subunit encoded by *ORF61,* and p56 an epitope derived from the viral single-stranded DNA biding protein encoded by *ORF6.* Transcripts from these genomic regions were detected in fibroblasts infected with RDV-50.stop, suggesting that these are RTA-independent genes which are expressed in abortively infected cells or transiently expressed during latency. Following WT MHV68 challenge, virus-specific CD8 T cells were further elevated by 7 days post-infection, suggesting that RDV-50.stop vaccination primed the T cell response for the challenge. Viral titers from lung homogenates collected on day 7 were significantly reduced by three orders of magnitude in the vaccinated mice, often to levels below the limit of detection. It is interesting to speculate that the T cell response that occurred after WT virus challenge in vaccinated animals reflects an increase in the local ability to control infection at the mucosal barrier. Future studies may evaluate whether the increase in virus-specific T cells are due to recruitment or expansion. Taken together, pre-existing humoral and cellular immunity in response to prime-boost immunization with a RDV controls acute replication. This response was durable as protection was maintained out to 3 months post-boost.

At 16 days post-infection in unvaccinated mice, acute infection has been cleared and latency is established. Indeed virus-specific T cell responses in sham-vaccinated mice surpassed those of vaccinated animals after WT virus challenge presumably due to the higher viral load present in the absence of immune control afforded by the vaccine. RDV-50.stop led to a 10-fold reduction in genome-positive splenocytes compared to sham-vaccinated control mice. Likewise, reactivation was nearly abolished within the vaccinated animals. However, the viral load of vaccinated animals based on the frequency of genome-positive cells appeared to recover to the levels of unvaccinated mice by 42 dpi. Considering the inability of cells harboring RDV-50.stop to undergo lytic replication and the strong block in reactivation that correlates with humoral and effector CD8 T cell responses in the vaccinated mice at 16 dpi, this recovery is likely due to proliferative expansion of latent cells. This finding highlights a major challenge for prophylactic vaccine efforts against the gammaherpesviruses.

Within immunocompromised individuals, GHV infection can lead to the onset of a plethora of lymphoproliferative disorders^68^. Previously studied latency-defective, yet replication-competent viruses were found to provide protection from WT MHV68 challenge^36,37^, but a replication-competent virus may pose a risk for hosts lacking innate immune control. Because of this, we wanted to determine if RDVs were both safe and effective in reducing the incidence of disease in a model of innate immune deficiency. To eliminate the risk of WT contamination and reversion, we grew RDV-50.stop on cell lines expressing a codon-shuffled *ORF50* construct, limiting sequence homology to inhibit recombination while still providing the WT protein required for replication^43^. Indeed, we previously reported that 1×10^6^ infectious RDV-50.stop particles produced on the codon-shuffled producer cells was not lethal in SCID mice, which occurs with as few as 10 PFU of WT virus^43^. To determine if RDV vaccination was protective in cases where a host is susceptible to severe disease, we vaccinated mice lacking the type I interferon receptor common alpha subunit*. Ifnar1^-/-^* mice experience acute disease leading to mortality following MHV68 infection^59,60^. RDV prime-boost vaccination of *Ifnar1^-/-^* mice increased virus-specific antibody titers with neutralizing capacity, similar to immune competent C57BL/6 mice. RDV vaccination led to a striking, complete protection from WT MHV68 challenge, as all vaccinated mice maintained consistent body weights through infection and displayed no outward signs of distress. Of those sham-vaccinated *Ifnar1^-/-^* mice that survived to the endpoint of the experiment, latency was reduced ten-fold compared to vaccinated controls. RDV vaccination is effective at inducing immune responses that correlate with protection from acute infection and disease in hosts susceptible to severe disease.

While our data demonstrate the protective potential of an RDV-based vaccine strategy, there are a few caveats with the RDV vaccine platform. While we noted lower overall levels of latent infection following WT GHV challenge, differential PCR demonstrates that RDV vaccination did not yield sterilizing immunity, as WT MHV68 comprises the bulk of the latent reservoir in vaccinated animals. It is worth noting, the phase 2 clinical trial of gp350 administration blocked a substantial degree of infectious mononucleosis even though EBV infection was not reduced^29,30^. Given the remarkable capacity of GHVs to promote proliferation of latently infected B cells in the absence of lytic replication, sterilizing immunity may be an unattainable goal. However, vaccination strategies that reduce EBV-driven IM and MS, and the numerous lymphomas and cancers caused by both EBV and KSHV is still a clinically significant outcome^9,31^. Another caveat of this first-generation design is that RDV-50.stop leaves intact numerous genes that enable the establishment and maintenance of latency, notably the unique MHV68 *M* genes and *ORF73* encoding LANA. As such, RDV-50.stop is able to establish a chronic latent infection, albeit at lower levels than WT virus at 16 dpi. Live-attenuated viruses that are unable to establish latency protect from WT MHV68 challenge^36,37^. A replication competent MHV68 vaccine virus lacking LANA, the latency protein essential for episomal maintenance, was reported to provide better protection than a replication-dead virus that lacked both the RTA lytic transactivator and LANA^37^. Such a contrast is not surprising since only single vaccinations in the absence of adjuvant were evaluated^37^. Here, the RDV-50.stop virus was administerd in the absence of adjuvant. While boosting enhanced the humoral response to infection, the CD8 T cell response was only moderately impacted. Future studies will evaluate whether combinatorial replication and latency-defective viruses benefit from adjuvanted boosting and reach the immunogenicity required for protection.

In summary, the studies described here detail the potential of a RDV-based vaccine platform to generate immune responses capable of inhibiting GHV replication, latency, and reactivation in immune competent mice and protection from death in a lethal model of disease. We observed virus-specific humoral and CD8 T cell responses upon RDV-RDV-50.stop vaccination; however, the full repertoire of the immune reactome remains to be defined. Rationally designed vaccines tested in the MHV68 model system will enable targeted depletion and transfer of specific immune components to delineate the determinants and effector mechanisms that block and control infection. Further refinement of vaccine design by delaying the block in virus replication to lytic genes of later kinetic classes and co-administration of next generation RDVs in the context of adjuvant is expected to further expand the breadth and depth of antigen presentation and immune correlates of protection.

## Methods

### Cells and viruses

Primary C57BL/6 murine embryonic fibroblasts (MEFs), NIH 3T12 (ATCC CCL-164), BHK21 (ATCC CCL-10), Vero (ATCC CCL-81), and Vero-Cre cells were cultured in Dulbecco’s Modified Eagle Medium (DMEM) supplemented with 10% fetal bovine serum (FBS), 100 ug/ml penicillin, 100 ug/ml streptomycin, and 2 mM L-glutamine to form complete DMEM (cDMEM). Cells were maintained at constant conditions in a standard incubator at 37°C, 5% CO_2,_ and 99% humidity. Primary MEFs at passages 2 and 3 were used for limiting dilution reactivation assays.

Viruses utilized in this study include WT MHV68 (WUMS strain) for vaccine challenge experiments in C57BL/6 mice, BAC-derived WT MHV68^69^ for comparative immune responses in C57BL/6 and vaccine challenge experiments in *Ifnar1^-/-^* mice, and the replication-dead (RDV) ORF50.stop recombinant MHV68 produced on codon-shuffled RTA-expressing NIH3T12 cells as previously described^41^ for which JCF and LTK hold a patent [US Patent 11,149,255].

### RNA-sequencing

NIH 3T12 fibroblasts were infected with RDV-50.stop or WT MHV68 at MOI 3 for 18 hours. Samples were harvested in Trizol, chloroform-extracted, and then RNA was isolated with the Qiagen RNeasy kit including the on-column DNase I treatment. Library preparation and sequencing were conducted at Azenta Life Sciences (South Plainfield, NJ, USA).

Extracted RNA samples were quantified using Qubit 2.0 Fluorometer (Life Technologies, Carlsbad, CA, USA) and RNA integrity was checked using Agilent TapeStation 4200 (Agilent Technologies, Palo Alto, CA, USA).

RNA sequencing libraries were prepared using the NEBNext Ultra II RNA Library Prep Kit for Illumina following manufacturer’s instructions (NEB, Ipswich, MA, USA). Briefly, mRNAs were first enriched with Oligo(dT) beads. Enriched mRNAs were fragmented at 94 °C. First strand and second strand cDNAs were subsequently synthesized. cDNA fragments were end repaired and adenylated at 3’ends, and universal adapters were ligated to cDNA fragments, followed by index addition and library enrichment by limited-cycle PCR. The sequencing libraries were validated on the Agilent TapeStation (Agilent Technologies, Palo Alto, CA, USA), and quantified by using Qubit 2.0 Fluorometer (Invitrogen, Carlsbad, CA) as well as by quantitative PCR (KAPA Biosystems, Wilmington, MA, USA).

RNA-seq data were aligned and counted using the CCR Collaborative Bioinformatics Resource (CCBR) in-house pipeline (https://github.com/CCBR/Pipeliner). Briefly, reads were trimmed of low-quality bases and adapter sequences were removed using Cutadapt v1.18 (http://gensoft.pasteur.fr/docs/cutadapt/1.18). Mapping of reads to custom reference hybrid genome described below was performed using STAR v2.7.0f in 2-pass mode^70,71^. Then, RSEM v1.3.0 was used to quantify gene-level expression ^72^ with quantile normalization and differential expression of genes analysis performed using limma-voom v3.38.3^73^. The data discussed in this publication have been deposited in NCBI’s Gene Expression Omnibus and are accessible through GEO Series accession GSExxxxx.

The custom reference genome allowing quantification of both viral and host expression used in this alignment consisted of the mouse reference genome (mm10/Apr. 2019/GRCm38) FASTA with a MHV68 FASTA sequence added as an additional pseudochromosome (Supplementary File “MHV68_Krug.fa”). This viral genome was prepared from the annotated herpesvirus genome (NCBI reference) with the addition of a CMV-driven Histone H2B-YFP fusion protein locus found in our mutant MHV68 virus strain, utilized to track individually infected cells. The custom gene annotations used for gene expression quantification consisted of a concatenation of the mm10 GENCODE annotation version M21^74^ and annotations of the MHV68 genome. All overlapping regions of the viral ORFs were removed to create a minimal, non-overlapping annotation. The sequencing library used was unstranded, making stranded counts of highly-overlapping viral ORFs problematic. This annotation was therefore used to make conservative estimates of the expression of individual viral genes. The custom viral GTF annotation files used for this quantification is provided as Supplemental File “M21_MHV68_Krug_Nonoverlapping.gtf”.

Data visualizations for RNAseq were created using Prism (GraphPad), and R (https://www.R-project.org/)^75^, RStudio (http://www.rstudio.com/)^76^, and the pheatmap package (https://CRAN.R-project.org/package=pheatmap)^77^. For both visualizations, counts were normalized first for library size, counts per million mapped reads (CPM), then for composition bias, trimmed-mean of M-values (TMM) before import to RStudio where they were log_2_ transformed prior to heatmap creation using the pheatmap package. For the purposes of visualization, +1 was added to each CPM TMM value prior to log_2_ transformation.

### Animal studies

All animal protocols that were performed by Stony Brook University staff were approved by the Institutional Animal Care and Use Committee of Stony Brook University. All animal procedures reported in this study that were performed by NCI-CCR affiliated staff were approved by the NCI Animal Care and Use Committee (ACUC) and in accordance with federal regulatory requirements and standards. All components of the intramural NIH ACU program are accredited by AAALAC International. Mouse experiments were carried out in accordance with guidelines from the National Institute of Health, UAMS Division of Laboratory Animal Medicine (DLAM), and the UAMS Institutional Animal Care and Use Committee (IACUC). The protocols supporting these animal studies were approved by the UAMS IACUC prior to the beginning of the study. Mice were anesthetized prior to inoculation. Mice were assessed twice daily for signs of disease and distress and measured once each day to monitor body weight as an indicator of disease progression. Mice were humanly euthanized at the indicated experimental endpoints or upon displaying signs of distress, characterized by displaying lethargy, dehydration, or a body weight reduction of 20% or more from the initial measurement taken on the day of inoculation.

Male and female C57BL/6 mice were purchased from Harlan/Envigo RMS (Indianapolis, IN) or Jackson Laboratories (Bar Harbor, ME). Male and female *Ifnar1^-/-^* (B6.(Cg)-*Ifnar11tm1.2Ees*/J) mice were ordered from Jackson Laboratories (Bar Harbor, ME). Eight to ten-week-old mice were anesthetized using an isoflurane chamber and inoculated with 10^6^ PFU of RDV-50.stop diluted in 200 uL DMEM intraperitoneally and boosted with an equivalent dose on days 25 and 39 post-priming. Mock vaccinated and vaccinated C57BL/6 mice were challenged by IN infection with 1×10^3^ PFU of WT MHV68 diluted in 20 μl complete DMEM. *Ifnar1^-/-^* mice were challenged with 2×10^6^ PFU WT MHV68 a dose shown previously to be lethal in approximately 50% of cases^60^. Serum and splenocytes were collected by submandibular vein sampling, or following humane euthanasia or at the endpoint of the study.

### Flow cytometry

2 × 10^6^ single cell suspensions prepared from the lung or spleen were resuspended in 200 ul of PBS with 2% fetal bovine serum and blocked with TruStain fcX (BioLegend, San Diego, CA). T cell subsets were identified with antibodies against CD45 (clone 30-F11, BV510), TCR; (clone H57-597, PerCP-Cy5.5), CD8 (clone 53-6.7, BV785), CD44 (clone IM7, APC-eFluor780), CD62L (clone MEL-14, FITC), CD127 (clone 678 A7R34, PE-Dazzle594), and KLRG1 (clone 2F1/KLRG1, PE-Cy7). All antibodies were purchased from BioLegend, eBioscence (Thermo Fisher Scientific), or BD Biosciences (San Jose, CA). H-2K(b)-p79 or H-2D(b)-p56 MHC-peptide complex was provided as dextramers conjugated to PE or APC (Immundex, Fairfax, VA).The data was collected on a CytoFLEX flow cytometer (Beckman Coulter) and analyzed using FlowJoX v10.0.7 (Treestar Inc., Ashland, OR). Cells were first gated as live per exclusion of Alexa Fluor™ 700 NHS Ester uptake and singlet lymphocytes based on forward and side scatter parameters prior to subgating.

### Peptide stimulation

For analysis of T cell effector responses, 1 × 10^6^ splenocytes were plated into each well of a 96-well flat-bottom plate and either left untreated or treated with 1 ug/ml p56 (AGPHNDMEI) or p79 (TSINFVKI) peptides (Genscript, Piscataway, NJ) for 5 h at 37°C in the presence of Brefeldin A (BD Cytofix/Cytoperm, BD Biosciences). The Fc receptors were blocked prior to surface staining with antibodies against CD45, TCRbeta, CD8, CD4 (clone GK1.5, BV711), CD44, Upon fixation and permeabilization with the BD Cytofix/Cytoperm kit (BD Biosciences), cells were stained with antibodies to IFNγ (clone XMG1.2, PE-Cy7) and TNFα (clone MP6-XT22, BV650).

### Pathogenesis assays

For acute titers, mice were euthanized with isoflurane at seven days post-infection, and both lungs were removed and frozen at −80°C. Lungs were disrupted in 1 ml of 8% cMEM using 1 mm zirconia beads in a bead beater (Biospec, Bartlesville, OK) and plated on NIH 3T12 cells. NIH 3T12 cells were plated at a density of 1.8 x 10^5^ cells/mL one day prior to infection. Serial dilutions of cell homogenate were added to the cell monolayer for 1 hr at 37°C, with rocking every 15 minutes, followed by an overlay of 5% methylcellulose (Sigma) in cMEM and incubated at 37°C. After 7-8 days, cells were fixed with 100% methanol (Sigma) and stained with 0.1% crystal violet (Sigma) in 20% methanol, and plaques were scored.

For limiting dilution analysis, spleens were homogenized in a tenBroek tissue disrupter. Red blood cells were lysed by incubating in 8.3 g/L ammonium chloride for 10 minutes at room temperature with constant shaking. RBC lysis was neutralized with 25 mL DMEM. Cells were filtered through a 40-micron filter to reduce clumping for further analyses. Limiting-dilution (LD)-PCR analyses to quantify frequencies of latently infected splenocytes were performed as previously described^61^. Briefly, 3-fold serial dilutions of latently infected cells were diluted in a background of uninfected 3T12 fibroblasts. After overnight digestion with proteinase K, cells were subjected to a nested PCR targeting the ORF50 region of the viral genome. Single-copy sensitivity and the absence of false-positive amplicons were confirmed using control standards. Amplicons were visualized for quantitation using ethidium bromide staining in 1.5% agarose gel electrophoresis.

*Ex vivo* reactivation efficiency was determined as previously described^78^. Briefly, 2-fold serial dilutions of latently infected splenocytes were plated on MEF or BHK21 monolayers for analysis of reactivation from splenocytes from C57BL/6 or *Ifnar1^-/-^*, respectively. The presence of preformed infectious virus was detected by plating mechanically disrupted cells on indicator monolayers in parallel. Cytopathic effect was scored 14 and 21 d after plating on MEFs or 6 to 7 d post-infection on BHK21.

### Immunoblot

3T12 fibroblasts were mock-infected or infected with WT MHV68. Lysates were diluted in Laemmli sample buffer and resolved by sodium dodecyl sulfate-polyacrylamide gel electrophoresis (SDS-PAGE). Resolved proteins were transferred to nitrocellulose membranes (Thermo Scientific), and blots were probed with mouse serum harvested from sham and vaccinated animals diluted 1:1000 in a 1% milk in TBS with 0.1% Tween-20 (TBS-T). Antibody-bound proteins were recognized using horseradish peroxidase (HRP)-conjugated goat anti-mouse IgG (Jackson ImmunoResearch). A chemiluminescent signal was detected using a ChemiDoc MP imaging system (Bio-Rad) on blots treated with Clarity ECL reagent (Bio-Rad).

### Neutralization assay

Neutralization was tested by means of a plaque reduction neutralization test (PRNT) as described^79^. Briefly, three-fold serum dilutions, starting from an initial concentration of 1:80 in cDMEM were incubated with 150 PFU of MHV68 on ice for one hour. The virus/serum mixture was then added to a sub-confluent BHK21 monolayer (4 × 10^4^ cells/well) plated the previous day in a 24-well plate, in triplicate. As a control, three wells received no-serum added virus. Infected cells were overlaid with methylcellulose and incubated at 37°C for four days. Methylcellulose media was then aspirated, and cell monolayers were stained with a solution of crystal violet (0.1%) in formalin. Percent neutralization was determined by comparison of the number of plaques in experimental wells compared to no-serum added control wells, and each data point was the average of three wells.

### Enyzme-linked immunosorbent assays (ELISA)

To measure antibody-specificity of serum, high binding plates (MidSci) were coated with 0.5% paraformaldehyde-fixed viral antigens in carbonate buffer (0.0875 M Na_2_CO_3_, 0.0123 M HCO_3_, pH 9.2) and incubated overnight at 4°C. Plates were washed in PBS with 0.05% Tween-20 (PBS-T) and blocked with 5% powdered milk-PBS-T for 1 hour at 37°C. Mouse sera were serially diluted in 1% milk-PBS-T and incubated for 1 hour at 37°C. IgG was detected by incubation with HRP-conjugated donkey anti-mouse IgG (Jackson ImmunoResearcg). SureBlue substrate (KPL) was added to detect virus-specific antibodies and read at 450 nm on a BioTek Synergy LX plate reader.

### PCR genotyping

Nucleic acid was isolated from splenocytes at the conclusion of the study. Total DNA was isolated using a GenCatch blood and tissue genomic mini-prep kit (Epoch Life Science). A nested PCR for the detection of the FRT-scar present in RDV-50.stop MHV68 was performed using 200 ng of genomic DNA with GoTaq polymerase (Promega) using primers (dPCR_R1_for GGACCACGCTTTCCAGAGAA, dPCR_R1_rev TCTGGTGGGATGTTGATGGC, dPCR_R2_for CCATGTGGGTACATCTAGCTTC, and dPCR_R2_rev CCAACACATTGCGCCCAAATGTC). Cycling conditions were as previously described^61^. In parallel, a nested PCR for the detection of both WT and RDV-50.stop MHV68 was performed using 200 ng of genomic DNA with GoTaq polymerase and ORF50 primers as previously described^61^. Amplicons were visualized for quantitation. Frequency of RDV-50.stop MHV68 infection was reported as the percentage of FRT-scar amplicons divided by the total MHV68 ORF50 amplicons.

### Statistical Analysis

All data was analyzed using GraphPad Prism software (GraphPad Software, http://www.graphpad.com<u>, La Jolla, CA</u>). Frequencies of immune cells analyzed by one-way or two-way ANOVA followed by post-tests depending on parametric distribution. Total numbers of an immune cells subset per animal were log_10_-transformed prior to ANOVA. Titer data were analyzed with unpaired t-test or one-way ANOVA for multiple groups. Based on the Poisson distribution, the frequencies of viral genome–positive cells and reactivation were obtained from the nonlinear regression fit of the data where the regression line intersected 63.2%. Extrapolations were used for samples that did not intersect 63.2%. The log_10_-transformed frequencies of genome-positive cells and reactivation were analyzed by unpaired, two-tailed t-test or one-way ANOVA for multiple groups.

## Acknowledgements

We thank Dr. Tiffany Weinkopff at UAMS for providing lodging to W.A.B. during a winter storm, enabling completion of lethal challenge experiments. This work utilized the computational resources of the NIH HPC Biowulf cluster (http://hpc.nih.gov).

This research was supported in part by the Intramural Research Programs of the NIH, the National Cancer Institute (NCI) (L.T.K.) S.M.O. was supported by a postdoctoral fellowship from Translational Research Institute (TRI) grant TL1 TR003109 through the NIH National Center for Advancing Translational Sciences. J.C.F. was supported by NIH NCI R01CA167065 and NIH NIAID R21AI139580. C.K. and B.S.S. were supported by The G. Harold and Leila Y. Mathers Foundation grant MF-1901-00210 (B.S.S.).

## Author Contributions

J.C.F. and L.T.K. conceived the study. W.A.B., S.O., and C.K. designed the study. W.A.B., K.M., L.B., V.K., and L.T.K. performed animal experimentation. W.A.B., S.O., K.M., performed sample processing in the laboratory. W.A.B., S.O., K.M., C.H.H., L.B., V.K., C.K., B.S.S., J.C.F. and L.T.K. analyzed the data. W.A.B, S.O., C.H.H., J.C.F, and L.T.K. wrote the article. All authors reviewed the article and agreed to its contents.

