## Supplemental Text and Figures for "Replication-dead gammaherpesvirus vaccine protects against acute replication, reactivation from latency, and lethal challenge in mice"

Supplemental File “M21_MHV68_Krug_Nonoverlapping.gtf”

### Supplemental Materials and Methods

To determine the sensitivity of primers that targeted the FRT locus, we performed a nested PCR reaction on RDV-50.stop BAC DNA using both our designed FRT-targeted primers (dPCR_R1_for GGACCACGCTTTCCAGAGAA, dPCR_R1_rev TCTGGTGGGATGTTGATGGC [805 bp product in RDV-50.stop]; dPCR_R2_for CCATGTGGGTACATCTAGCTTC and dPCR_R2_rev CCAACACATTGCGCCCAAATGTC [308 bp product]), as well as traditional pan-MHV68 primers (KM86 AACTGGAACTCTTCTGTGGC and KM89 GGCCGCAGACATTTAATGAC [586 bp product]; KM87 CCCCAATGGTTCATAAGTGG and KM88 ATCAGCACGCCATCAACATC [382 bp product])[61]. BAC DNA was serially diluted from $10^7$ copies per reaction to $10^{-3}$ copies per well. Cycling conditions were as previously described[61]. Amplicons were visualized for quantitation. Frequencies of amplification were comparable between FRT-targeted and pan-MHV68 primer pairs.

Nucleic acid was isolated from splenocytes at the conclusion of the study. Total DNA was isolated using a GenCatch blood and tissue genomic mini-prep kit (Epoch Life Science). A nested PCR for the detection of the FRT-scar present in RDV-50.stop MHV68 was performed using 200 ng of genomic DNA with GoTaq polymerase (Promega) using FRT-targeted primers. Cycling conditions were as previously described[61]. In parallel, a nested PCR for the detection of both WT and RDV-50.stop MHV68 was performed using 200 ng of genomic DNA with GoTaq polymerase and pan-MHV68 primers as previously described[61]. Amplicons were visualized for quantitation. Frequency of RDV-50.stop MHV68 infection was reported as the percentage FRT-scar amplicons of the total pan-MHV68 amplicons.

Supplemental Figure Legends

**Supplemental Fig 1 Virus-specific CD8 T cell responses upon a prime-boost regimen in C57BL/6 mice.** Virus p56-specific effector CD8-T cell response based on p56 tetramer+ of $CD44^{hi}CD62L^{-}$ CD8 T cells 14 d post prime-boost with RDV-50.stop. Naïve mice were age-matched, non-vaccinated controls. (Left) Representative gating strategy from naïve mice or mice infected with RDV-50.stop of WT virus. (Right) Total p56-reactive CD8 T cells per spleen of individual mice after initial prime and sequential boosts. Symbols represent individual mice (N=3-5) for boost 1 and bars are mean values; *, p<0.05; ***, p<0.0002; ****, p<0.0001 in Tukey's multiple comparisons post-text of two-way ANOVA.

**Supplemental Fig 2 RDV-50.stop establishes latency and does not induce sterilizing immunity.** C57BL/6 mice were either sham-vaccinated or vaccinated and then boosted with $1x10^6$ PFU RDV-50.stop followed by challenge with $1x10^3$ PFU WT MHV68 at 16 d post-boost. **a** The frequency of latency determined by limiting dilution nested PCR of intact splenocytes for the viral genome at 42 d post-challenge of mice (N=3) with one or two boosts post-prime. **b** PCR genotyping of a pool of splenocytes from the indicated sets of mice (N=4-5) mock at 16 d post-challenge. To differentiate the RDV-50.stop vaccine virus from WT challenge virus, nested PCR was performed with primers that target the FRT sequence only present within RDV-50.stop, in parallel with 'pan-MHV68' primers that detect both RDV-50.stop and WT MHV68. For each set, bars indicate the percentage of PCR reactions that produced RDV-50.stop FRT amplimers as a percentage of reactions that produced pan-MHV68 amplimers. The absence of RDV-50.stop FRT amplimers in samples that yielded pan-MHV68 amplimers was considered WT.

**Supplemental Fig 3 Evaluation of immune response to MHV68 at seven days post-challenge with WT virus.** C57BL/6 mice were either sham-vaccinated or vaccinated twice (prime+boost) with $1x10^6$ PFU RDV-50.stop followed by mock challenge or challenge with $1x10^3$

PFU WT MHV68 at 15 d post-boost and analyzed 7 d post-challenge. **a** Total number of $CD44^{hi}$ CD8 T cells in the spleen that were reactive with the viral p56 epitope. **b** p56-tetramer+ CD8 T cells were further analyzed for markers of short-lived effector cell (SLEC, $KLRG^{+}CD127^{-}$) and **c** memory precursor effector cell subsets (MPEC, $KLRG^{-}CD127^{+}$). **d** MPECs were further delineated into $CD62L^{-}$ effector and $CD62L^{+}$ central MPECs. For **a-d**, symbols represent individual mice, (N=4-5); bars and whiskers are mean +/- SD values.*, p<0.05; ***, p<0.001; ****, p<0.0001 in Sidak's multiple comparisons test of one-way ANOVA between the indicated groups. **e** Virus-specific IgG from sham or RDV-50.stop vaccinated mice at 7 d post challenge measured by ELISA. **f** Virus neutralization in serum as determined by a plaque reduction assay. The $PRNT_{50}$ value is the dilution of serum to reach 50% neutralization of plaques. Symbols represent individual mice (N=3-5); bars are mean values +/- SD. *, p<0.05 in unpaired t test.

**Supplemental Fig 4 Evaluation of the T cell response to MHV68 p56 in vaccinated mice at 16 days post-challenge with WT virus.** C57BL/6 mice were either sham-vaccinated or vaccinated twice (prime+boost) with $1x10^6$ PFU RDV-50.stop MHV68 followed by challenge with $1x10^3$ PFU WT MHV68 at 15 days post-boost. **a** Total p56-tetramer+ CD8 T cells per spleen of individual mice after initial prime and sequential boosts. Percentage of p56-tetramer+ CD8 T cells with markers of **b** short-lived effector cell (SLEC, $KLRG^{+}CD127^{-}$) and **c** memory precursor effector cell subsets (MPEC, $KLRG^{-}CD127^{+}$). **d** MPECs were further delineated into $CD62L^{-}$ effector and $CD62L^{+}$ central MPECs for p79- and p56-tetramer+ CD8 T cells. For each graph, symbols represent individual mice, (N=3-5); bars and whiskers are mean +/- SD values. **, p<0.05; ***, p<0.001; ****, p<0.0001 in Sidak's multiple comparisons test of one-way ANOVA between the indicated groups.

**Supplemental Fig 5 Vaccination with RDV-50.stop leads to durable protection against wild-type MHV68 challenge.** C57BL/6 mice were either sham-vaccinated or vaccinated twice (prime+boost) with $1x10^6$ PFU RDV-50.stop followed by challenge with $1x10^3$ PFU WT MHV68 at 90 days post-boost. **a-b** Total p79- or p56-tetramer+ CD8 T cells per lung. **c,g** Total p79- or p56-tetramer+ CD8 T cells per lung. Tetramer+ CD8 T cells were further analyzed for markers of **d,h** short-lived effector cell (SLEC, $KLRG^+CD127^-$) and **e,i** memory precursor effector cell subsets (MPEC, $KLRG^-CD127^+$). **f,j** MPECs were further delineated into CD62L- effector and CD62L+ central MPECs. **k** Percentage of $CD44^{hi}$ CD8 T cells producing both TNF$\alpha$ and IFN$\gamma$ after peptide stimulation. For each graph, symbols represent individual mice, (N=4-5); bars and whiskers are mean +/- SD values.*, $p<0.05$; **, $p<0.05$; ***, $p<0.001$; ****, $p<0.0001$ in Sidak's multiple comparisons test of one-way ANOVA between the indicated groups.

**Supplemental Fig 6 *Ifnar1*$^{-/-}$ mice generate neutralizing humoral immune responses following RDV-50.stop vaccination.** *Ifnar1*$^{-/-}$ mice were immunized with $1x10^6$ PFU of RDV-50.stop in the peritoneum. Mice were given two booster doses of $1x10^6$ PFU RDV-50.stop and serum was collected on day 52 post-vaccination. **a** Virus-specific IgG from sham or RDV-50.stop vaccinated mice measured by ELISA. **b** Virus neutralization in serum as determined by plaque assay. The $PRNT_{50}$ value is the dilution of serum to reach 50% neutralization of plaques. Symbols represent pooled serum run in technical triplicate. Error bars represent standard error of the means. **, $p<0.005$ as determined by an unpaired, two-tailed student's t-test.

**Supplemental Fig 7 Vaccination with RDV-50.stop MHV68 protects mice susceptible to severe disease from weight-loss.** *Ifnar1*$^{-/-}$ mice (n=10) were either sham-vaccinated or vaccinated with 3 doses of $1x10^6$ PFU RDV-50.stop MHV68 and challenged with a lethal dose of $2x10^6$ PFU WT MHV68. (A) Sham-vaccinated and (B) vaccinated mice were weighed daily for 42 days to measure disease progression. Symbols represent individual mice; error bars represent standard error of the means.

**Supplemental Fig 8 RDV-50.stop establishes latency and does not induce sterilizing immunity in *Ifnar1$^{-/-}$* mice.** *Ifnar1$^{-/-}$* mice were either sham-vaccinated or vaccinated with 3 doses of $1x10^6$ PFU RDV-50.stop and challenged with a lethal dose of $2x10^5$ PFU WT MHV68. **a** The frequency of latency determined by limiting dilution nested PCR of intact splenocytes for the viral genome at 20 d post-challenge. **b** PCR genotyping of a pool of splenocytes from RDV-50.stop vaccinated and challenged mice (N=10) at 20 d post-challenge. To differentiate the RDV-50.stop vaccine virus from WT challenge virus, nested PCR was performed with primers that target the FRT sequence only present within RDV-50.stop, in parallel with 'pan-MHV68' primers that detect both RDV-50.stop and WT MHV68. For each set, bars indicate the percentage of PCR reactions that produced RDV-50.stop FRT amplimers as a percentage of reactions that produced pan-MHV68 amplimers. The absence of RDV-50.stop FRT amplimers in samples that yielded pan-MHV68 amplimers was considered WT.

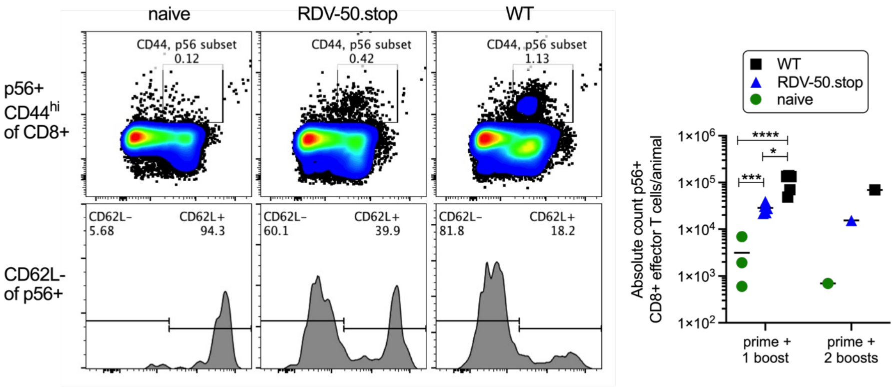


**Supplemental Fig 1 Virus-specific CD8 T cell responses upon a prime-boost regimen in C57BL/6 mice.** Virus p56-specific effector CD8 T cell response based on p56 tetramer+ of CD44$^{hi}$CD62L$^{-}$ CD8 T cells 14 d post prime-boost with RDV-50.stop. Naïve mice were age-matched, non-vaccinated controls. (Left) Representative gating strategy from naïve mice or mice infected with RDV-50.stop of WT virus. (Right) Total p56-reactive CD8 T cells per spleen of individual mice after initial prime and sequential boosts. Symbols represent individual mice (N=3-5) for boost 1 and bars are mean values; *, p<0.05; ***, p<0.0002; ****, p<0.0001 in Tukey's multiple comparisons post-text of two-way ANOVA.

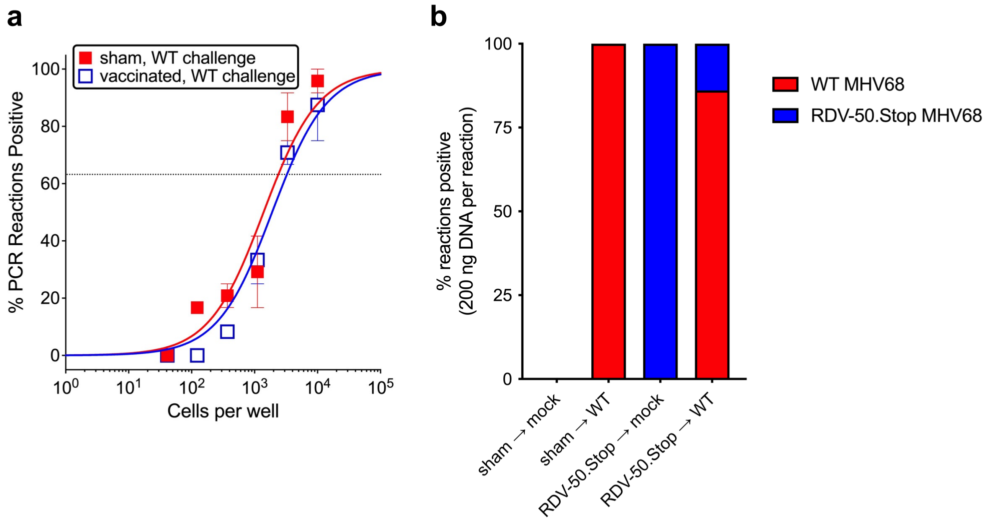


**Supplemental Fig 2 RDV-50.stop establishes latency and does not induce sterilizing immunity.** C57BL/6 mice were either sham-vaccinated or vaccinated and then boosted with 1x$10^6$ PFU RDV-50.stop followed by challenge with 1x$10^3$ PFU WT MHV68 at 16 d post-boost. **a** The frequency of latency determined by limiting dilution nested PCR of intact splenocytes for the viral genome at 42 d post-challenge of mice (N=3) with one or two boosts post-prime. **b** PCR genotyping of a pool of splenocytes from the indicated sets of mice (N=4-5) mock at 16 d post-challenge. To differentiate the RDV-50.stop vaccine virus from WT challenge virus, nested PCR was performed with primers that target the FRT sequence only present within RDV-50.stop, in parallel with 'pan-MHV68' primers that detect both RDV-50.stop and WT MHV68. For each set, bars indicate the percentage of PCR reactions that produced RDV-50.stop FRT amplimers as a percentage of reactions that produced pan-MHV68 amplimers. The absence of RDV-50.stop FRT amplimers in samples that yielded pan-MHV68 amplimers was considered WT.

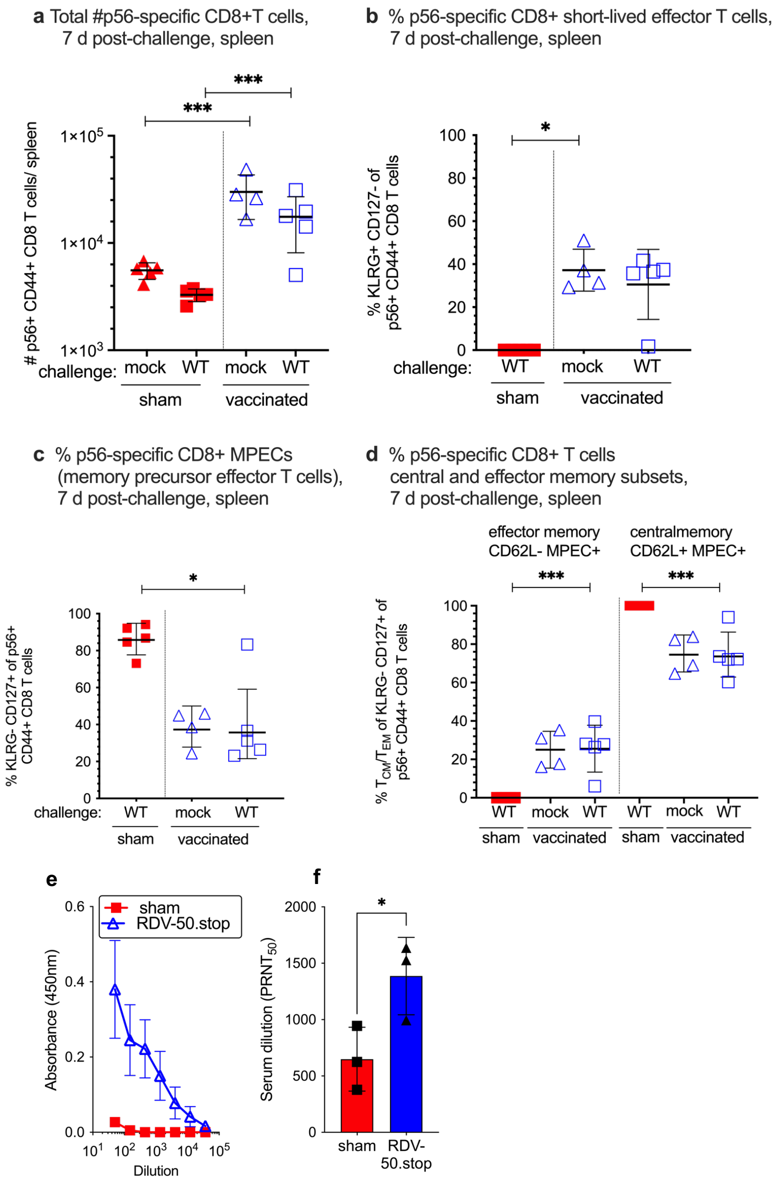


**Supplemental Fig 3 Evaluation of immune response to MHV68 at seven days post-challenge with WT virus.** C57BL/6 mice were either sham-vaccinated or vaccinated twice (prime+boost) with 1x10$^{6}$ PFU RDV-50.stop followed by mock challenge or challenge with 1x10$^{3}$ PFU WT MHV68 at 15 d post-boost and analyzed 7 d post-challenge. **a** Total number of CD44$^{hi}$ CD8 T cells in the spleen that were reactive with the viral p56 epitope. **b** p56-tetramer+ CD8 T cells were further analyzed for markers of short-lived effector cell (SLEC, KLRG$^{+}$CD127$^{-}$) and **c** memory precursor effector cell subsets (MPEC, KLRG$^{-}$CD127$^{+}$). **d** MPECs were further delineated into CD62L$^{-}$ effector and CD62L$^{+}$ central MPECs. For **a-d**, symbols represent individual mice, (N=4-5); bars and whiskers are mean +/- SD values.*, p<0.05; ***, p<0.001; ****, p<0.0001 in Sidak's multiple comparisons test of one-way ANOVA between the indicated groups. **e** Virus-specific IgG from sham or RDV-50.stop vaccinated mice at 7 d post challenge measured by ELISA. **f** Virus neutralization in serum as determined by a plaque reduction assay. The PRNT$_{50}$ value is the dilution of serum to reach 50% neutralization of plaques. Symbols represent individual mice (N=3-5); bars are mean values +/- SD. *, p<0.05 in unpaired t test.

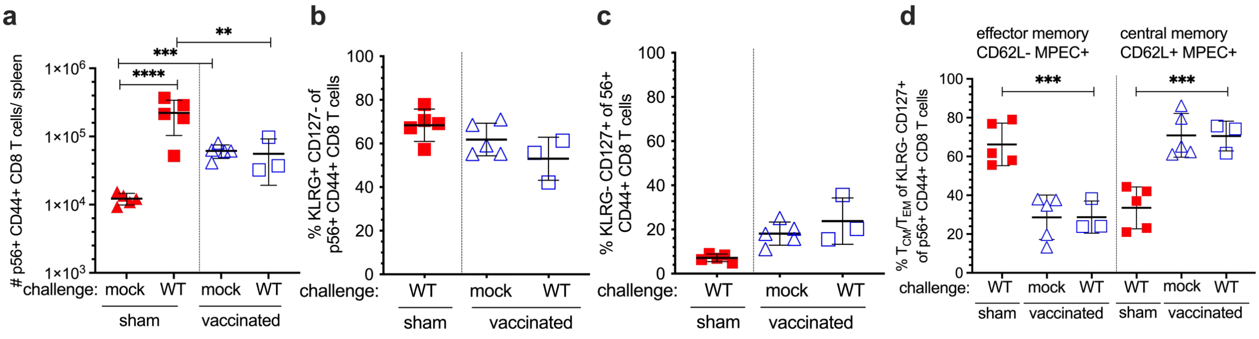


**Supplemental Fig 4 Evaluation of the T cell response to MHV68 p56 in vaccinated mice at 16 days post-challenge with WT virus.** C57BL/6 mice were either sham-vaccinated or vaccinated twice (prime+boost) with $1x10^6$ PFU RDV-50.stop MHV68 followed by challenge with $1x10^3$ PFU WT MHV68 at 15 days post-boost. **a** Total p56-tetramer+ CD8 T cells per spleen of individual mice after initial prime and sequential boosts. Percentage of p56-tetramer+ CD8 T cells with markers of **b** short-lived effector cell (SLEC, $KLRG^+CD127^-$) and **c** memory precursor effector cell subsets (MPEC, $KLRG^-CD127^+$). **d** MPECs were further delineated into $CD62L^-$ effector and $CD62L^+$ central MPECs for p79- and p56-tetramer+ CD8 T cells. For each graph, symbols represent individual mice, (N=3-5); bars and whiskers are mean +/- SD values. **, p<0.05; ***, p<0.001; ****, p<0.0001 in Sidak's multiple comparisons test of one-way ANOVA between the indicated groups.

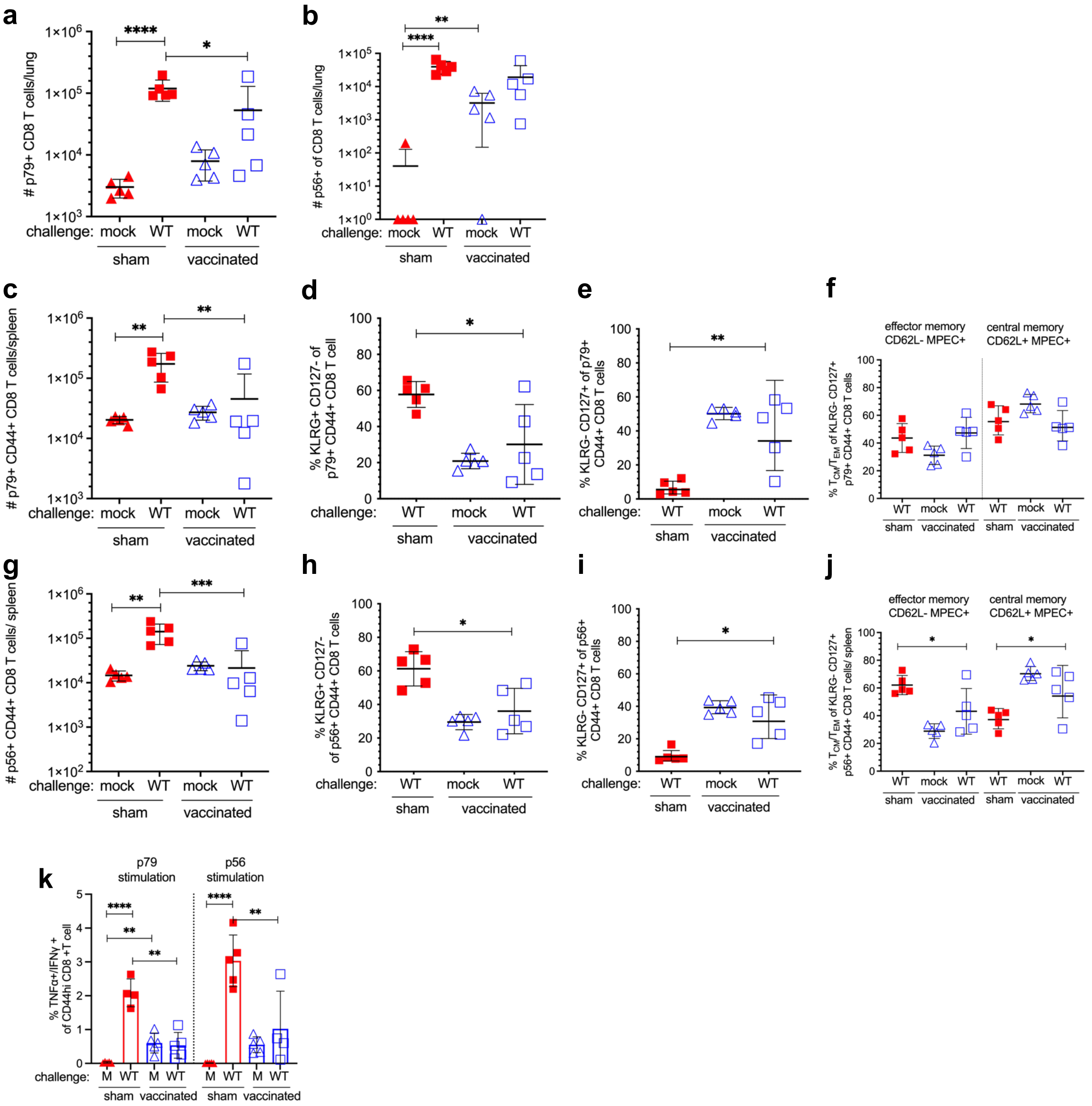


**Supplemental Fig 5 Vaccination with RDV-50.stop leads to durable protection against wild-type MHV68 challenge.** C57BL/6 mice were either sham-vaccinated or vaccinated twice (prime+boost) with 1x$10^6$ PFU RDV-50.stop followed by challenge with 1x$10^3$ PFU WT MHV68 at 90 days post-boost. **a-b** Total p79- or p56-tetramer+ CD8 T cells per lung. **c,g** Total p79- or p56-tetramer+ CD8 T cells per lung. Tetramer+ CD8 T cells were further analyzed for markers of **d,h** short-lived effector cell (SLEC, $KLRG^{+}CD127^{-}$) and **e,i** memory precursor effector cell subsets (MPEC, $KLRG^{-}CD127^{+}$). **f,j** MPECs were further delineated into CD62L- effector and CD62L+ central MPECs. **k** Percentage of $CD44^{hi}$ CD8 T cells producing both TNFα and IFNγ after peptide stimulation. For each graph, symbols represent individual mice, (N=4-5); bars and whiskers are mean +/- SD values.*, $p<0.05$; **, $p<0.05$; ***, $p<0.001$; ****, $p<0.0001$ in Sidak's multiple comparisons test of one-way ANOVA between the indicated groups.

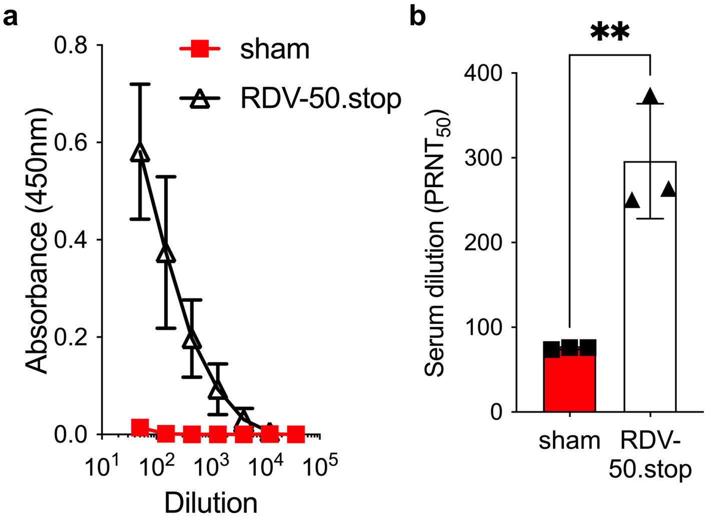


**Supplemental Fig 6** ***Ifnar1**$^{-/-}$* **mice generate neutralizing humoral immune responses following RDV-50.stop vaccination.** *Ifnar1*$^{-/-}$ mice were immunized with 1x10$^6$ PFU of RDV-50.stop in the peritoneum. Mice were given two booster doses of 1x10$^6$ PFU RDV-50.stop and serum was collected on day 52 post-vaccination. **a** Virus-specific IgG from sham or RDV-50.stop vaccinated mice measured by ELISA. **b** Virus neutralization in serum as determined by plaque assay. The $PRNT_{50}$ value is the dilution of serum to reach 50% neutralization of plaques. Symbols represent pooled serum run in technical triplicate. Error bars represent standard error of the means. **, $p<0.005$ as determined by an unpaired, two-tailed student's t-test.

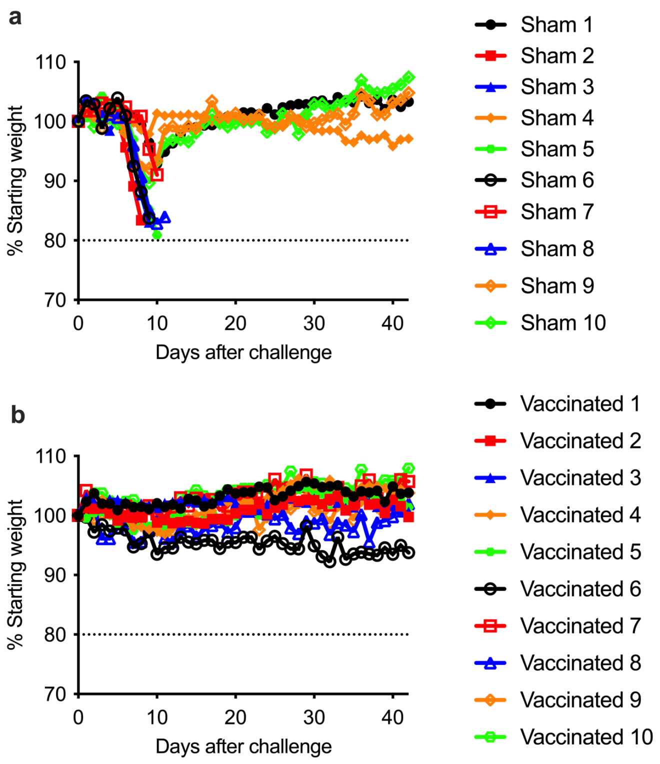


**Supplemental Fig 7 Vaccination with RDV-50.stop MHV68 protects mice susceptible to severe disease from weight-loss.** *Ifnar1$^{-/-}$* mice (n=10) were either sham-vaccinated or vaccinated with 3 doses of $1x10^6$ PFU RDV-50.stop MHV68 and challenged with a lethal dose of $2x10^6$ PFU WT MHV68. (A) Sham-vaccinated and (B) vaccinated mice were weighed daily for 42 days to measure disease progression. Symbols represent individual mice; error bars represent standard error of the means.

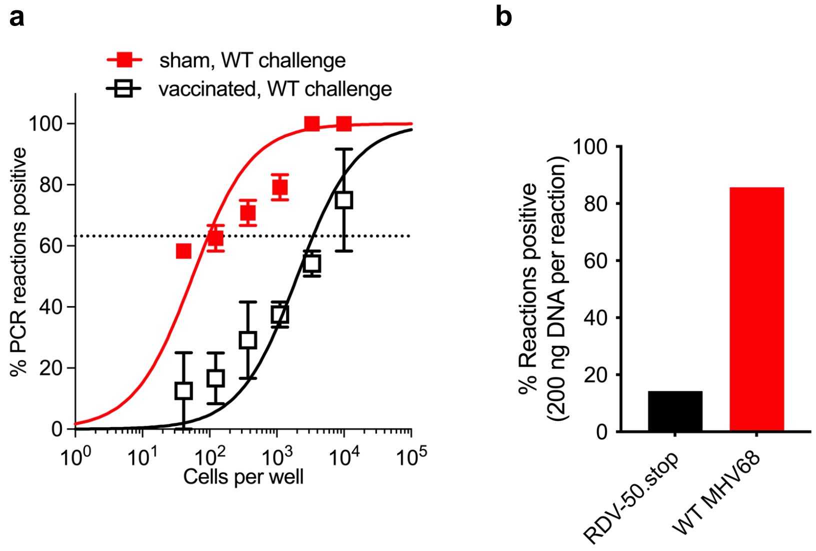


**Supplemental Fig 8 RDV-50.stop establishes latency and does not induce sterilizing immunity in *Ifnar1$^{-/-}$* mice.** *Ifnar1$^{-/-}$* mice were either sham-vaccinated or vaccinated with 3 doses of $1x10^6$ PFU RDV-50.stop and challenged with a lethal dose of $2x10^5$ PFU WT MHV68. **a** The frequency of latency determined by limiting dilution nested PCR of intact splenocytes for the viral genome at 20 d post-challenge. **b** PCR genotyping of a pool of splenocytes from RDV-50.stop vaccinated and challenged mice (N=10) at 20 d post-challenge. To differentiate the RDV-50.stop vaccine virus from WT challenge virus, nested PCR was performed with primers that target the FRT sequence only present within RDV-50.stop, in parallel with 'pan-MHV68' primers that detect both RDV-50.stop and WT MHV68. For each set, bars indicate the percentage of PCR reactions that produced RDV-50.stop FRT amplimers as a percentage of reactions that produced pan-MHV68 amplimers. The absence of RDV-50.stop FRT amplimers in samples that yielded pan-MHV68 amplimers was considered WT.
